# HSP70 is upregulated after heat but not freezing stress in the freeze-tolerant cricket *Gryllus veletis*

**DOI:** 10.1101/2024.10.30.621172

**Authors:** Victoria E. Adams, Maranda L. van Oirschot, Jantina Toxopeus

**Affiliations:** Department of Biology, St. Francis Xavier University, 2320 Notre Dame Ave, Antigonish NS, Canada, B2G 2W5

**Keywords:** cold tolerance, insect, molecular chaperone, protein denaturation, thermal physiology

## Abstract

Heat shock proteins (HSPs) are well known to prevent and repair protein damage caused by various abiotic stressors, but their role in low temperature and freezing stress is not well-characterized compared to other thermal challenges. Ice formation in and around cells is hypothesized to cause protein damage, yet many species of insects can survive freezing, suggesting HSPs may be an important mechanism in freeze tolerance. Here, we studied HSP70 in a freeze-tolerant cricket *Gryllus veletis* to better understand the role of HSPs in this phenomenon. We measured expression of one heat-inducible HSP70 isoform at the mRNA level (using RT-qPCR), as well as the relative abundance of total HSP70 protein (using semi-quantitative Western blotting), in five tissues from crickets exposed to a survivable heat treatment (2 h at 40°C), a 6-week fall-like acclimation that induces freeze tolerance, and a survivable freezing treatment (1.5 h at -8°C). While HSP70 expression was upregulated by heat at the mRNA or protein level in all tissues studied (fat body, Malphigian tubules, midgut, femur muscle, nervous system ganglia), no tissue exhibited HSP70 upregulation within 2 – 24 h following a survivable freezing stress. During fall-like acclimation to mild low temperatures, we only saw moderate upregulation of HSP70 at the protein level in muscle, and at the RNA level in fat body and nervous tissue. Although HSP70 is important for responding to a wide range of stressors, our work suggests that this chaperone may be less critical in the preparation for, and response to, moderate freezing stress.

**Highlights:**

- Heat shock protein 70 (HSP70) may not contribute substantially to freeze tolerance
- Heat stress caused HSP70 mRNA and protein upregulation in the spring field cricket
- Acclimation prior to freezing was correlated with slight HSP70 upregulation
- HSP70 was not upregulated after freezing in this freeze-tolerant insect
- Further work is needed to determine whether freezing causes protein damage

**Graphical abstract:** 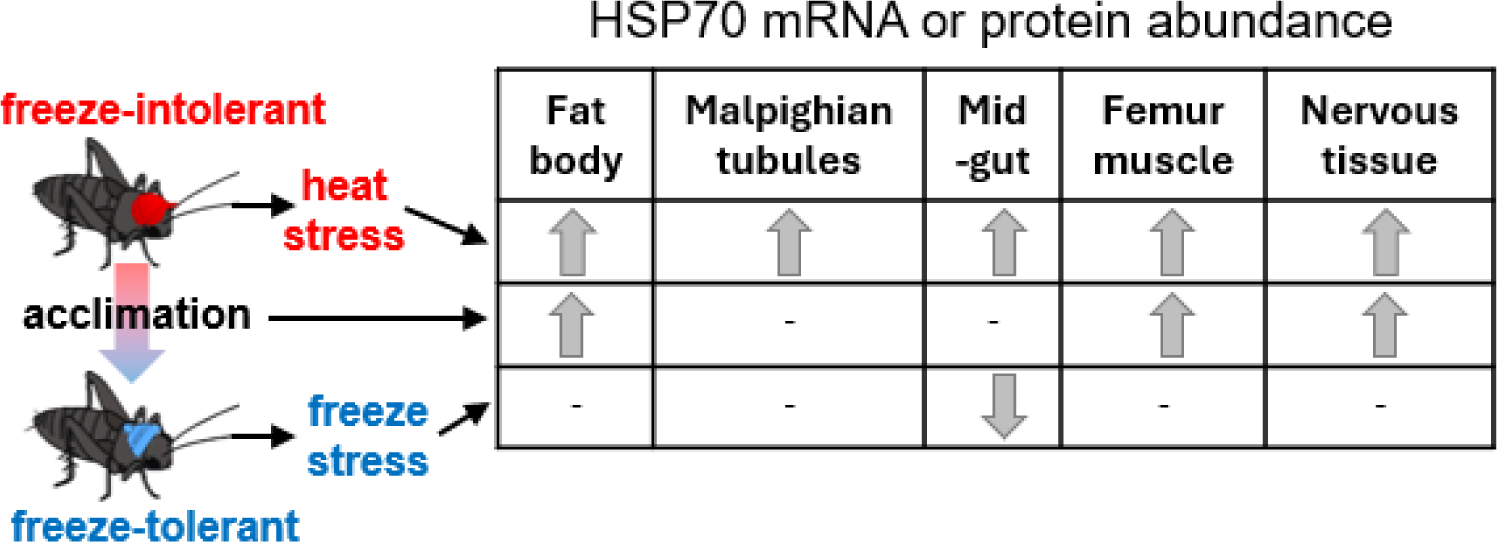

## Introduction

Many organisms overwinter in environments cold enough to freeze their body fluids, and this freezing is hypothesized to cause protein damage (Rozsypal, 2022; Toxopeus and Sinclair, 2018). Molecular chaperones such as heat shock proteins (HSPs) are important for preventing and repairing damage to proteins (Storey and Storey, 2023; Ullah et al., 2024), and are therefore likely candidates for helping organisms survive freezing (Teets et al., 2023; Toxopeus and Sinclair, 2018). Indeed, some freeze-tolerant insects upregulate heat shock protein 70 (HSP70) prior to (Des Marteaux et al., 2019) or after (Zhang et al., 2011) freezing stress, which is hypothesized to help them survive internal ice formation. Insects from more extreme cold environments such as Antarctica maintain constitutively high expression of HSP70 (Lencioni et al., 2015; Lopez-Martinez et al., 2008; Teets et al., 2020). However, recent work has shown that freezing of insects does not impair enzyme activities, even when the organism itself does not survive freezing (Grgac et al., 2022). This suggests that protein structure is not damaged by the freeze-thaw process (see Rozsypal, 2022), and therefore that HSPs may not be an important mechanism for supporting freeze tolerance. Here, we test the hypothesis that HSP70 is important for tolerating freezing and other thermal treatments in the freeze-tolerant spring field cricket *Gryllus veletis*.

Heat shock proteins are well-conserved molecular chaperones that are important for the cellular response to stress, as well as regular cellular processes involving the processing of proteins. HSPs can facilitate protein transport, folding, unfolding, assembly, and disassembly; degradation of misfolded proteins; and prevention of protein aggregation. HSPs can be classified based on molecular weight and function into categories such as small HSPs, HSP40, HSP60, HSP70, and HSP90, with the number referring to the protein mass in kDa (Feder and Hofmann, 1999; King and MacRae, 2015). Each class of HSP may have multiple isoforms within a species (Yocum, 2001), with patterns of gene expression that differ depending on the type of stress (Wang et al., 2019), life stage (Belén Arias et al., 2011), and tissue (King and MacRae, 2015). 70 kDa molecular chaperones that are constitutively expressed and involved in regular cellular protein maintenance can be referred to as heat shock cognates (HSC), distinguishing them from HSPs that are induced by stress and more important for stabilizing unfolded proteins and preventing aggregation (Clark and Worland, 2008; King and MacRae, 2015). HSP70s are particularly prevalent in studies of stress tolerance (Storey and Storey, 2023; Ullah et al., 2024), but have been examined in relatively few freeze-tolerant organisms.

*Gryllus veletis* (Orthoptera: Gryllidae) is distributed through much of southern Canada and the northern U.S.A., and overwinters as a freeze-tolerant fifth instar nymph (Alexander and Bigelow, 1960; Toxopeus et al., 2019c). In the laboratory, these nymphs undergo multiple physiological and biochemical adjustments to become freeze-tolerant following exposure to a 6-week fall-like acclimation regime of decreasing temperature and photoperiod (Toxopeus et al., 2019c, 2019b, 2019a). Previous transcriptomic work on fat body of these crickets identified one putative HSP70 as the only HSP upregulated during acclimation (Toxopeus et al., 2019a). However, transcriptomic studies alone are insufficient to confirm a role for HSPs in freeze tolerance. In this study, we first describe the sequence, structure, and expression of a putative HSP70 in response to heat shock in five *G. veletis* tissues to confirm its function as a heat-inducible HSP. We then evaluate the role of HSP70 in freeze tolerance by measuring HSP70 expression at the mRNA and protein level – again in five tissues – during fall-like acclimation and following a moderate freeze-treatment. While HSP70 is indeed heat-inducible in *G. veletis*, it is not upregulated following freezing stress, prompting additional hypotheses for further study.

## Methods

### Characterization of a putative Hsp70 sequence

Six putative transcripts from the *Gryllus veletis* transcriptome (Toxopeus et al., 2019a) had high similarity to other orthopteran (locust) HSP70 sequences (Table S1), and were therefore potential targets for studying gene expression. From this dataset, transcript Gvel_34771 (NCBI accession GGSD01051771.1) was the only sequence with a full open reading frame (ORF) that could encode a putative HSP70 protein (see Supplementary Information for more detail). We used Expasy’s Translate Tool (Gasteiger, 2003; https://web.expasy.org/translate/) to determine possible amino acid sequences from Gvel_34771. The amino acid and nucleotide sequences corresponding to the longest ORF aligned well with other known HSP70 and HSC70 proteins (Figures S1, S2), so we proceeded to assess its properties. We used SWISS-MODEL (Gasteiger, 2003; https://swissmodel.expasy.org/) to determine the predicted structure of our putative HSP70 polypeptide using default settings, and identified classic HSP70 domains by comparing our sequence to the domains described in Vostakolaei et al. (2021).

### Experimental design

We conducted three experiments to examine the impact of temperature on HSP70 expression at the mRNA and protein level in five tissues from *G. veletis* (Figure 1). To test if the crickets upregulated HSP70 during the acclimation that induces freeze tolerance, we compared crickets that were acclimated for three weeks or six weeks of fall-like conditions to control crickets that were never acclimated (Figure 1). To test if the HSP70 was heat-inducible, we compared crickets that were heat shocked and allowed to recover for 2 or 24 h to control crickets that were never acclimated or heat shocked (Figure 1). To test if the freeze-tolerant crickets upregulated HSP70 after freezing stress, crickets that were acclimated (freeze-tolerant), frozen, and allowed to recover for 2 or 24 h were compared to control crickets that were acclimated but never frozen (Figure 1). Fat body, Malpighian tubules, midgut, muscle (hind femur), and nervous tissue (ganglia) were dissected for each experimental group. There were 9 biological replicates for each treatment and tissue type (405 samples). Each biological replicate contained tissues pooled from three crickets.

**Figure 1.**
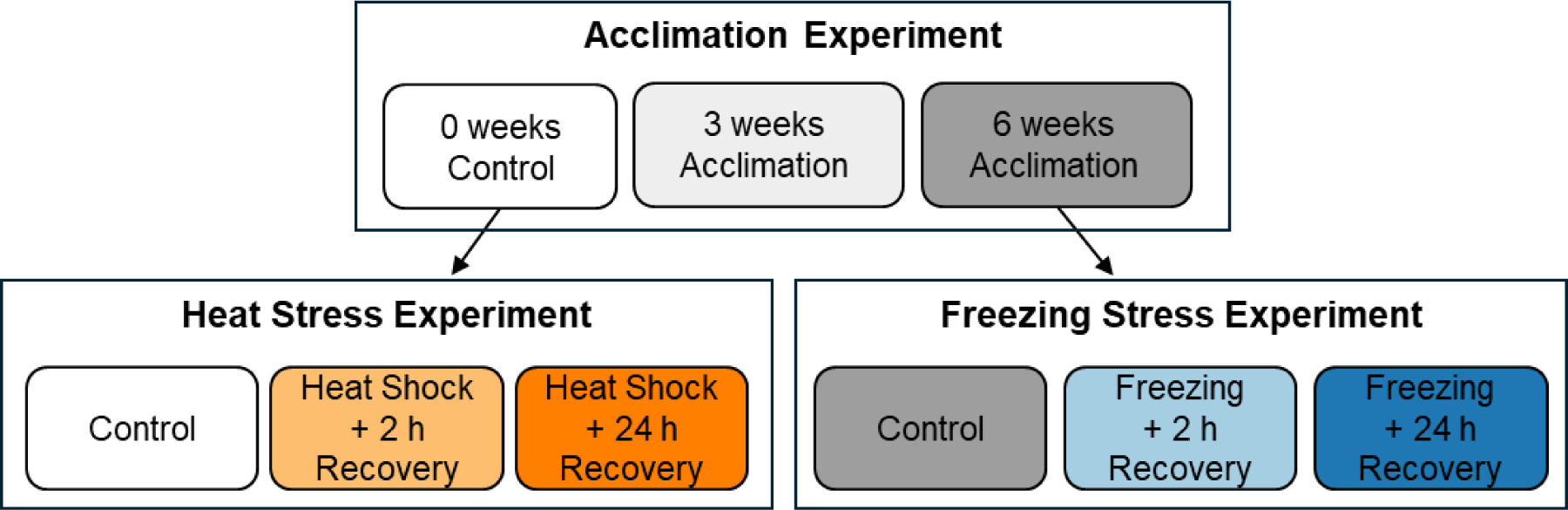
Experimental design for testing the effects of three thermal regimes on relative abundance of HSP70 mRNA and protein in *Gryllus veletis*. Each experiment contained three treatments: a control group, and two other groups that either differed in acclimation duration or the length of recovery following a thermal stress. Freeze-intolerant (0 weeks acclimation) crickets were used in the heat stress experiment, and freeze-tolerant crickets (6 weeks acclimation) were used in the freezing stress experiment.

### Insect rearing, acclimation, and temperature exposures

The spring field crickets used for these experiments were bred from a colony in the Animal Care facility at St. Francis Xavier University (Antigonish, NS, Canada), and obtained from an original source population in Lethbridge, AB, Canada (Toxopeus et al., 2019c). The crickets were reared as previously described (Toxopeus et al., 2019c) under summer-like conditions (25 °C, 14:10 Light: Dark photoperiod, 70% relative humidity). All experiments were conducted on fifth instar male nymphs. Nymphs were taken directly from rearing for use as the “0 week” group in the acclimation experiment, as well as for all groups in the heat stress experiment (Figure 1). To induce freeze tolerance, fifth instar male nymphs crickets were randomly selected and transferred to incubators (MIR-154-PA; PHCbi, Wood Dale, IL, USA) that mimicked fall-like conditions for 6 weeks (Table 1), with slight difference from the protocol described by Toxopeus et al., (2019c). Crickets were removed after 3 or 6 weeks of acclimation for use as the remaining groups of the acclimation experiment, and 6-week acclimated (freeze-tolerant) crickets were used for all groups in the freezing stress experiment (Figure 1).

**Table 1.** Acclimation that induced freeze tolerance in male 5^th^ instar *Gryllus veletis.* To mimic fall conditions, daily high temperatures (while lights were on) and low temperatures (while lights were off) decreased over 6 weeks, along with decreasing photoperiod.

| Week | Day | Day (lights on) |  | Night (lights off) |  |
| --- | --- | --- | --- | --- | --- |
|  |  | Duration (hours:minutes) | Temperature (°C) | Duration (hours:minutes) | Temperature (°C) |
| 1 | 1-3 | 11:30 | 16 | 12:30 | 12 |
|  | 4-7 | 11:12 | 15 | 12:48 | 11 |
| 2 | 8-10 | 10:54 | 14 | 13:06 | 9 |
|  | 11-14 | 10:36 | 12 | 13:24 | 8 |
| 3 | 15-17 | 10:18 | 11 | 13:42 | 7 |
|  | 18-21<sup>a</sup> | 10:00 | 9 | 14:00 | 6 |
| 4 | 22-24 | 9:42 | 8 | 14:18 | 4 |
|  | 25-28 | 9:24 | 6 | 14:36 | 3 |
| 5 | 29-31 | 9:06 | 4 | 14:54 | 2 |
|  | 32-35 | 8:48 | 3 | 15:12 | 2 |
| 6 | 36-38 | 8:30 | 2 | 15:30 | 1 |
|  | 39-42<sup>b</sup> | 8:12 | 1 | 15:48 | 0 |

<sup>a</sup>3 week acclimated crickets (21 days) and <sup>b</sup>6 week acclimated crickets (42 days) were collected/sampled at the end of the respective weeks.

Heat and freezing stress exposures were all conducted using previously-described methods (Lemay et al., 2024; McIntyre et al., 2023; Toxopeus et al., 2019c). Each cricket was placed into a 1.7 ml microcentrifuge tube, which we placed in a custom aluminum block that was temperature-controlled by a 1:1 propylene glycol:water mix circulated from an Arctic A25 Chiller (ThermoFisher, Mississauga, ON, Canada). Each cricket was in contact with a constantan-copper Type T thermocouple (Omega Engineering, Norwalk, CT, USA), which logged body temperature every second via a TC-08 USB unit interfaced with PicoLog 6.2.8 software (Pico Technology, Cambridgeshire, UK). After stress exposures, crickets were transferred to individual mesh-covered transparent cups containing rabbit food, water, and shelters made from egg cartons for recovery at room temperature.

Prior to the heat stress experiment (Figure 1), a pilot study was conducted on unacclimated crickets to determine an appropriate survivable heat shock temperature. Groups of 12 nymphs were exposed to one of five warm temperatures (38, 40, 42, 44, 46°C) for 2 h, followed by return to room temperature (c. 20°C) for recovery. Survival was assessed as previously described (Toxopeus et al., 2019c) two days post-heat shock. Based on these results, we chose a 2 h exposure at 40°C for the heat shock in our heat stress experiment, which is expected to result in 100% survival (see Results) but is likely stressful.

We conducted the freezing stress experiment using conditions that have resulted in high (92 ± 6%) survival of acclimated crickets previously (Toxopeus et al., 2019c). Each acclimated cricket had 2 µL of silver iodide slurry applied to its dorsal abdomen to help promote freezing (Toxopeus et al., 2019c). We gradually lowered the crickets’ body temperature at a rate of 0.25 °C/min from 4°C to -8°C, and confirmed freezing via the presence of a supercooling point (SCP; Sinclair et al., 2015). They were held at -8°C for 1.5 hours before being warmed to 4°C at a rate of 0.25°C/min, followed by recovery at room temperature.

### Tissue dissections

Five types of tissue were dissected from each cricket following its exposure to the conditions described in the experimental design (Figure 1). Dissections of the 0-week acclimation group and heat stressed crickets were performed at room temperature. Dissections of the 3-week and 6-week acclimation groups, as well as all freezing stress crickets were performed on ice. Tissues were flash frozen in 1.7 ml tubes in liquid nitrogen, and then stored at –80°C until RNA extractions or sample preparation for Western blotting.

### Measuring relative mRNA abundance with RT-qPCR

To prepare sample for RT-qPCR, RNA extractions were performed on each tissue sample using TRIzol reagent (300 µL) according to the manufacturer’s protocol (ThermoFisher) as previously described (Toxopeus et al., 2019a), and eluted in 30 µL of nuclease-free molecular biology grade water (Sigma-Aldrich, St. Louis, MO, USA). The quality and quantity of RNA was determined via absorbance at 230 nm, 260 nm, and 280 nm light using a NanoDropOne spectrophotometer. 0.2 µg of RNA was used for DNAse (ThermoFisher) treatment and cDNA synthesis via the iScript Reverse Transcription Supermix (BioRad, Mississauga, ON, Canada) according to the manufacturers’ protocols.

We designed primers to target a portion of the full HSP70 ORF represented by Gvel_34771 for use in RT-qPCR. The other five putative transcripts from (Toxopeus et al., 2019a) were all too short (< 750 bp; Table S1) to encode a full ORF, and therefore likely represented fragments of a full mRNA transcript or transcripts. Primers designed to target these other putative HSP70 transcript fragments (Table S1) did not produce amplicons (data not shown). We used Primer3 (https://primer3.ut.ee/) to design our primers (Table 2) and ordered them from ThermoFisher.

**Table 2.**
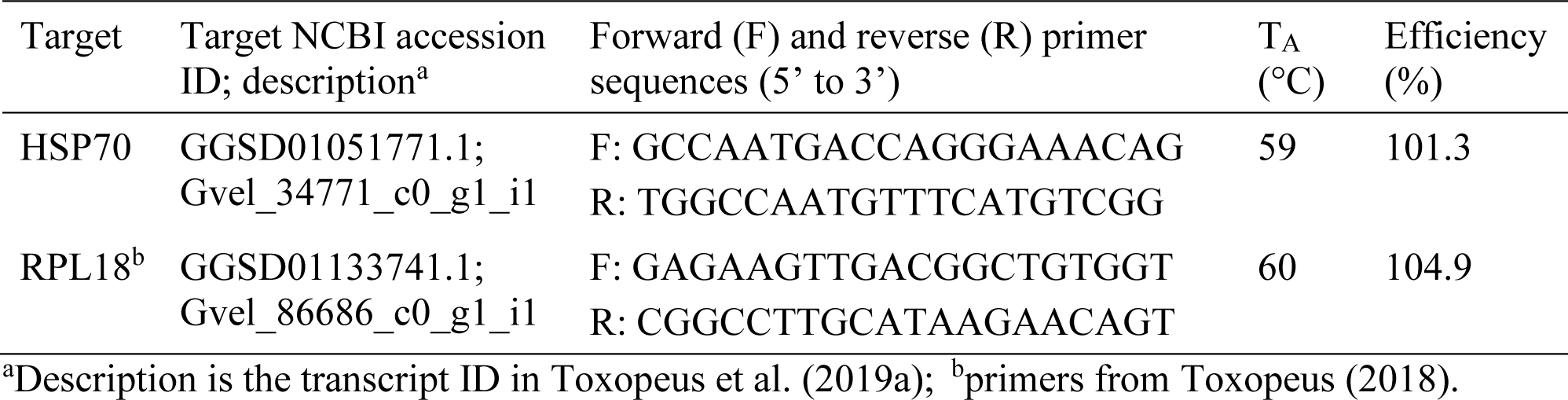
Primers used in this study. T_A_, annealing temperature used in PCR and RT-qPCR

Ribosomal Protein L18 (RPL18) was chosen as a reference gene (Table 2), based on previous success in *G. veletis* (Toxopeus, 2018). PCR amplicons were generated with each primer pair and *G. veletis* cDNA using DreamTaq (ThermoFisher) according to manufacturer’s instructions (Lemay et al., 2024; McIntyre et al., 2023). To verify that our RT-qPCR primers were specific for our target *G. veletis* HSP70 transcript (Gvel_34771) and RPL18, we sent the amplicons for Sanger sequencing at the Toronto Centre for Applied Genomics (Toronto, ON, Canada). Clustal Omega (Sievers et al., 2011; https://www.ebi.ac.uk/jdispatcher/msa/clustalo) was used to align our sequenced HSP70 amplicon to both Gvel_34771 and Gvel_19946 (NCBI Accession GGSD01027946.1), the latter of which overlaps with the region of Gvel_34771 that we were trying to amplify (Figure S3) but was not our intended target. RT-qPCR conditions described below were chosen because they ensured a primer efficiency between 90 and 110% (Table 2).

RT-qPCR was run in triplicate for each RNA sample described in the experimental design (Figure 1), using each of the RPL18 and HSP70 primers (Table 2). cDNA samples were diluted 1/10 prior to use in RT-qPCR for gene expression analysis. RT-qPCR was performed according to the manufacturer’s protocol (SsoAdvanced SYBR Green Supermix; BioRad) in 10 µL reaction mixes with 2 µL diluted cDNA, 400 nM primers and the following parameters: initial denaturation at 94°C for 5 min; 40 cycles of 94°C for 15 s, T_A_ (see Table 2) for 15 s, 72°C for 20 s; followed by melt curve determination via ramping from 65°C to 95°C in increments of 0.5°C. The 2^-ΔΔC^_T_ method (Livak and Schmittgen, 2001) was used to determine whether relative abundance of HSP70 mRNA changed in response to acclimation or thermal stress compared to respective control samples (Figure 1).

### Measuring relative protein abundance with Western blotting

To prepare tissue samples for Western blotting, 50-100 µL of potassium phosphate buffer (100 mM potassium phosphate, 0.1% Triton X, pH 6.9) was added to samples in 1.7 mL tubes, followed by homogenization with a plastic pestle. Homogenates were centrifuged for at 800 × *g* for 5 min at 4°C before collecting and storing supernatant at -80°C. Protein concentrations of samples were quantified using Pierce BCA Protein Assay Kit (ThermoFisher), followed by dilution with potassium phosphate buffer to the desired concentration. Samples were then diluted 1:1 with Laemmli sample buffer (65.8 mM Tris-HCl, pH 6.8, 2.1% SDS, 26.3% (w/v) glycerol, 0.01% bromophenol blue, 5% (v/v) 2-mercaptoethanol) and boiled in a water bath for 5 min to denature proteins.

For each tissue, an equal amount of protein was loaded per lane into an acrylamide gel (5% stacking, 10% resolving), and separated via SDS-PAGE alongside the IRIS11 Prestained Protein Ladder (FroggaBio, Toronto, ON, Canada), in a mini-PROTEAN Tetra System (BioRad) at 35 V for 30 min followed by 175 V for 45 min. Between 5 µg and 20 µg of protein was loaded on the gel, based on preliminary blots for each tissue that established the quantity of protein in control samples that resulted in target detection, but not saturation, of the chemiluminescence signal (King et al., 2013). Proteins were transferred onto a 0.45 µm nitrocellulose membrane (Cytiva Amersham Protran, ThermoFisher) using a semi-dry transfer at 25V for 30 min in the Trans-Blot Turbo transfer system (BioRad). To stain total protein, blots were washed 3 times for 5 min with distilled water and incubated in Ponceau stain (0.1% (w/v) Ponceau red dye in 5% acetic acid) for 5 min. The membrane was washed 3 times for 5 min in TBST (10 mM Tris, 140 mM NaCl, 0.1% Tween-20, pH 7.4) and imaged under white light for total protein quantification. The membrane was then incubated in a blocking solution of 4% bovine serum albumin (BSA) in TBST overnight (16-20 h) at 4°C, followed by incubation with 1:3,000 HSP70 monoclonal primary antibody (3A3; Fisher Scientific) in blocking solution for 1 h at room temperature, three 5 min washes in TBST, and incubation in the 1:3,000 blotting grade HRP-conjugated mouse secondary antibody (BioRad) in blocking solution for 30 min. Each blot was developed over 5 min with a Clarity Western ECL Substrate kit (BioRad), and imaged with a 20 s exposure in a ChemiDoc MP imaging system (BioRad).

The relative abundance of HSP70 protein in each sample compared to control samples (Figure 1) on the same blot was determined after correcting for any variation in total protein between lanes. Total protein abundance per sample was quantified from images of Ponceau-stained blots calculating the integrated density of each lane in Fiji (ImageJ software; Schindelin et al., 2012). HSP70 abundance per sample was quantified from chemiluminescence staining on each blot using the method outlined by Stael et al. (2022).

### Statistical analyses

For each experiment in our design (Figure 1), each tissue was analyzed separately, and statistical analyses were performed in R v4.4.1 (R Core Team, 2024). For each tissue, to determine whether the relative HSP70 mRNA abundance differed between treatment groups within an experiment, an ANOVA was performed on ΔΔCq values followed by a Tukey’s post-hoc test if the ANOVA was significant (α = 0.05). To test for differences in relative protein abundance in the thermal stress experiments, non-parametric Kruskal-Wallis tests and Dunn’s post-hoc tests were performed. For the acclimation experiment, relative protein abundance was analyzed via a Mann-Whitney test (non-parametric t-test) because only control (0 weeks acclimation) and fully acclimated (6 weeks acclimation) samples were available for Western blotting analysis.

## Results

### Characterization and specific amplification of putative HSP70 encoded by Gvel_34771

The predicted *Gryllus veletis* HSP70 protein based on Gvel_34771 was 659 amino acids long, and included the classic HSP70 domains (Figure 2). These included a nucleotide-binding domain (binds and hydrolyzes ATP) and a substrate-binding domain (Figure 2). The substrate binding domain consisted of a beta-pleated sheet region that can bind the substrate, and a C-terminal alphal-helical ‘lid’ (Figure 2).

**Figure 2.**
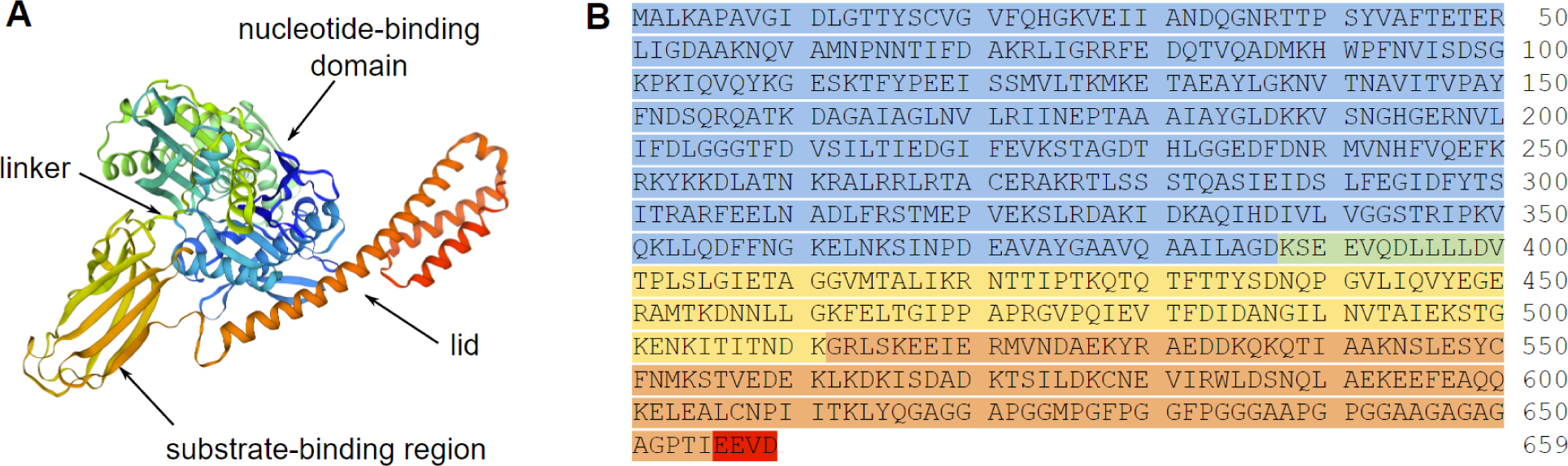
*Gryllus veletis* HSP70 (A) predicted structure and (B) sequence based on the longest open reading frame (ORF) from putative HSP70 transcript Gvel_34771. **(A)** The predicted structure was based on an HSC70 from *Melitaea cinxia* (UniProt accession T1SC35) in SWISS MODEL, and is colour-coded in a blue to red gradient from the N-terminus to the C-terminus. **(B)** The predicted amino acid sequence is colour-coded to identify the nucleotide-binding domain (blue), linker (green), substrate-binding region (yellow), lid (orange), and C-terminal EEVD domain (red), following Vostakolaei et al. (2021). The substrate-binding region and lid are both part of the substrate-binding domain.

### Putative HSP70 encoded by Gvel_34771 is heat-inducible

Unacclimated crickets showed moderate heat tolerance. All crickets survived brief (2 h) exposure to 38°C and 40°C, but survival declined sharply at temperatures above 42°C (Figure 3). In our heat stress experiment (Figure 1), 96% of crickets were alive at the time of dissection (2 or 24 h post-heat shock at 40°C), confirming that the heat stress was generally non-lethal.

**Figure 3.**
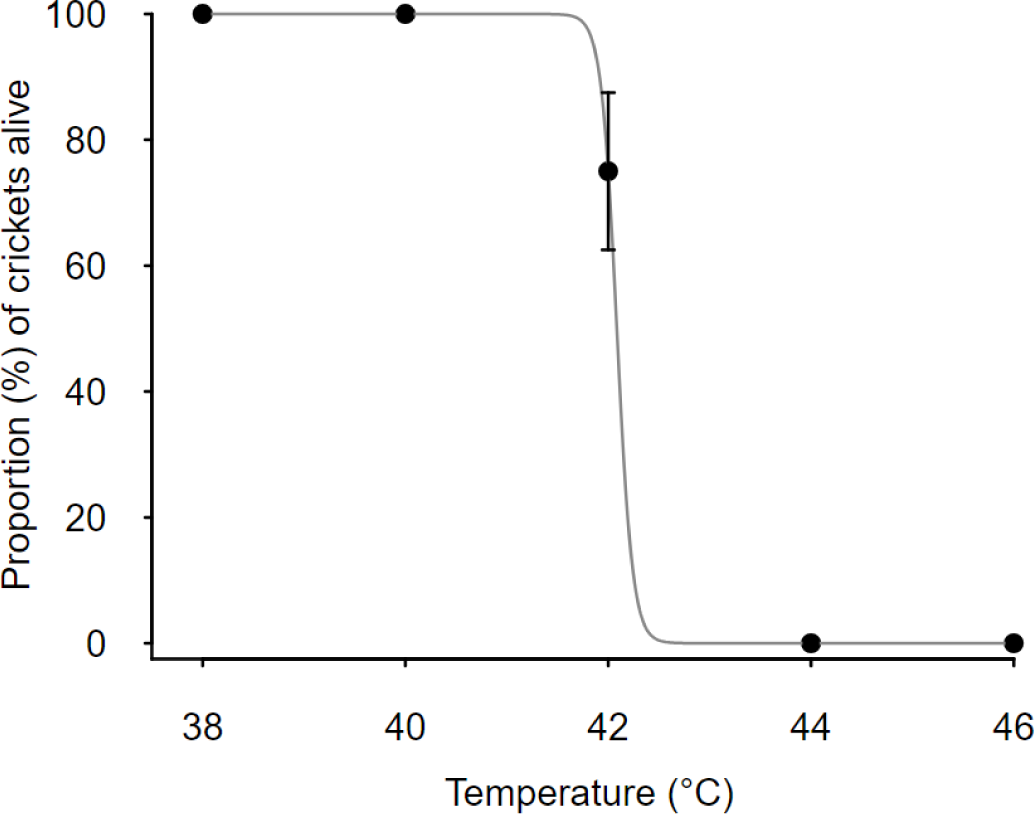
Proportion (%) of *Gryllus veletis* that survived following exposure to acute heat shock at the indicated temperatures. Temperature exposures were 2 h long. Each point represents a proportion (± standard error of proportion) determined from 12 crickets. Grey line indicates the equation derived from the logistic regression of survival data.

The primers we designed to measure relative expression of HSP70 mRNA were specific for their intended target, Gvel_34771. The HSP70 amplicon produced with our RT-qPCR primers had the expected size (Figure 4A) of c. 186 bp. The sequenced amplicon was an exact match to the targeted Gvel_34771 and had less than 80% identity with the non-targeted Gvel_19946 (Figure 4B), strongly suggesting that our primers specifically amplified the isoform of HSP70 encoded by Gvel_34771 alone. We therefore interpret the RT-qPCR results below as representative of one HSP70 isoform. The anti-HSP70 antibody we used for Western blotting resulted in detection of 70 kDa proteins in all five tissues we used in this study (Figure 4C). However, the anti-HSP70 antibody we used can recognize multiple HSP70 isoforms in multiple organisms (ThermoFisher), so we assume the relative protein abundance measurement in our study includes multiple HSP70 isoforms.

**Figure 4.**
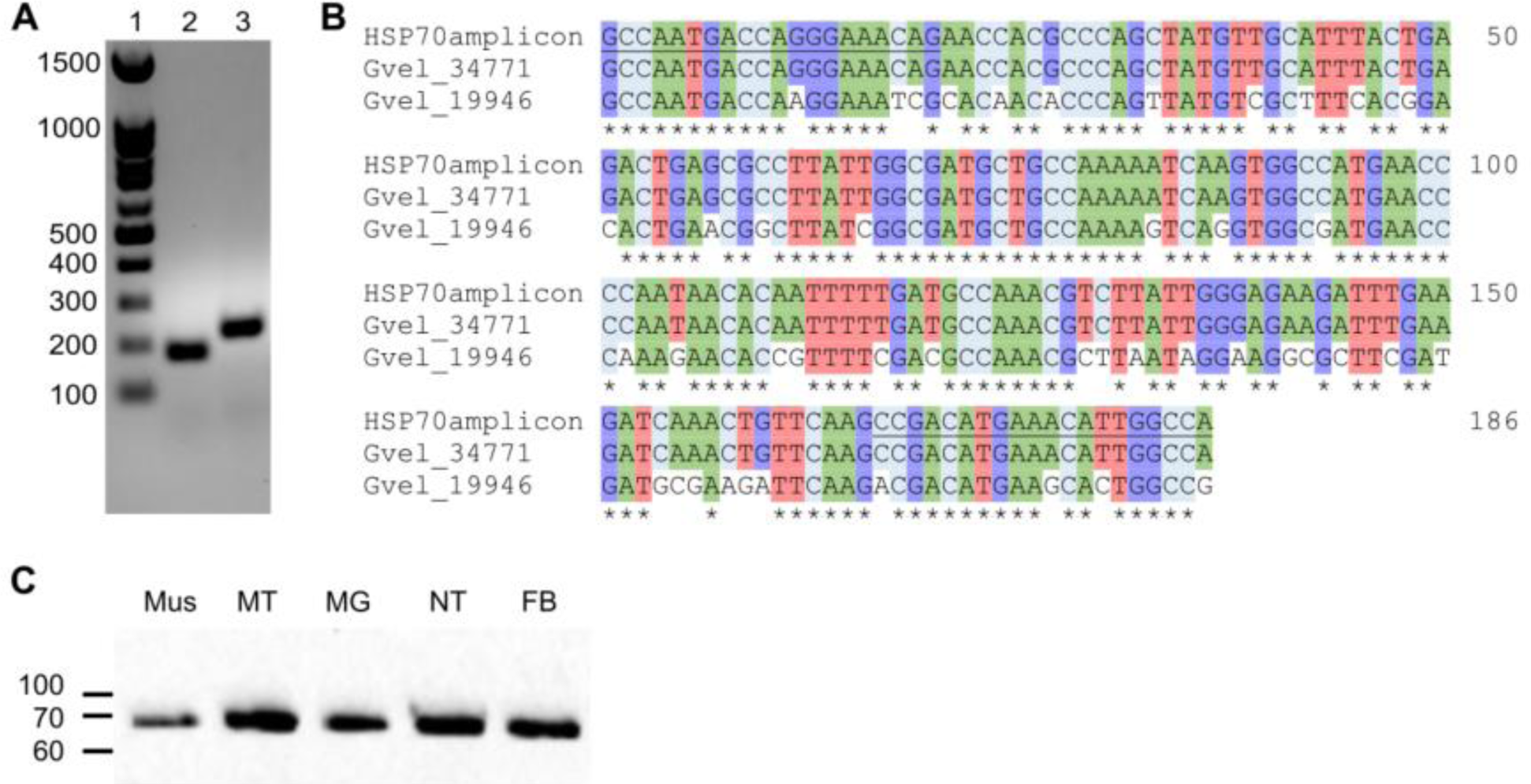
(A) Amplification and (B) sequence of the target HSP70 RT-qPCR product from *Gryllus veletis* cDNA, and (C) detection of HSP70 proteins from *G. veletis* tissues via Western blotting. **(A)** 1.5% agarose gel stained with RedSafe to show: 1. 100 bp ladder, 2. HSP70 amplicon, and 3. RPL18 amplicon (reference gene). Sizes of ladder fragments are indicated in base pairs (bp) to the left. **(B)** Alignment of the sequenced HSP70 amplicon with partial sequences of putative *Gryllus veletis* HSP70 transcripts Gvel_34771 (GGSD01051771.1; target) and Gvel_19946 (GGSD01027946.1; non-target). Forward and reverse primer sequences are underlined at the 5’ and 3’ ends of the amplicon. Nucleotides that match the amplicon sequence are colour-coded. **(C)** SDS-PAGE and immunostaining of HSP70 in 20 µg of protein extracted from five tissues. Sizes of ladder fragments are indicated in kiloDaltons (kDa). Mus, muscle (femur); MT, Malpighian tubules; MG, midgut; NT, nervous tissue (ganglia); FB, fat body.

The relative abundance of HSP70 mRNA and protein abundance significantly increased in at least one tissue after 2 or 24 h of recovery post-heat shock (Figure 5, Table 3), suggesting our target gene was heat inducible. At the mRNA level, HSP70 expression increased post-heat shock in Malpighian tubules, femur muscle tissue, and nervous tissue (Figure 5A). Conversely, in fat body and midgut HSP70 mRNA abundance decreased 24 h after heat shock compared to non-heat shocked controls (Figure 5A). In all tissues except nervous tissue, HSP70 protein abundance was increased by at least 50% post heat-shock (Figure 5B). A similar trend was present in nervous tissue, but relative HSP70 protein abundance was notably variable 24 h post heat-shock (Figure 5B).

**Figure 5.**
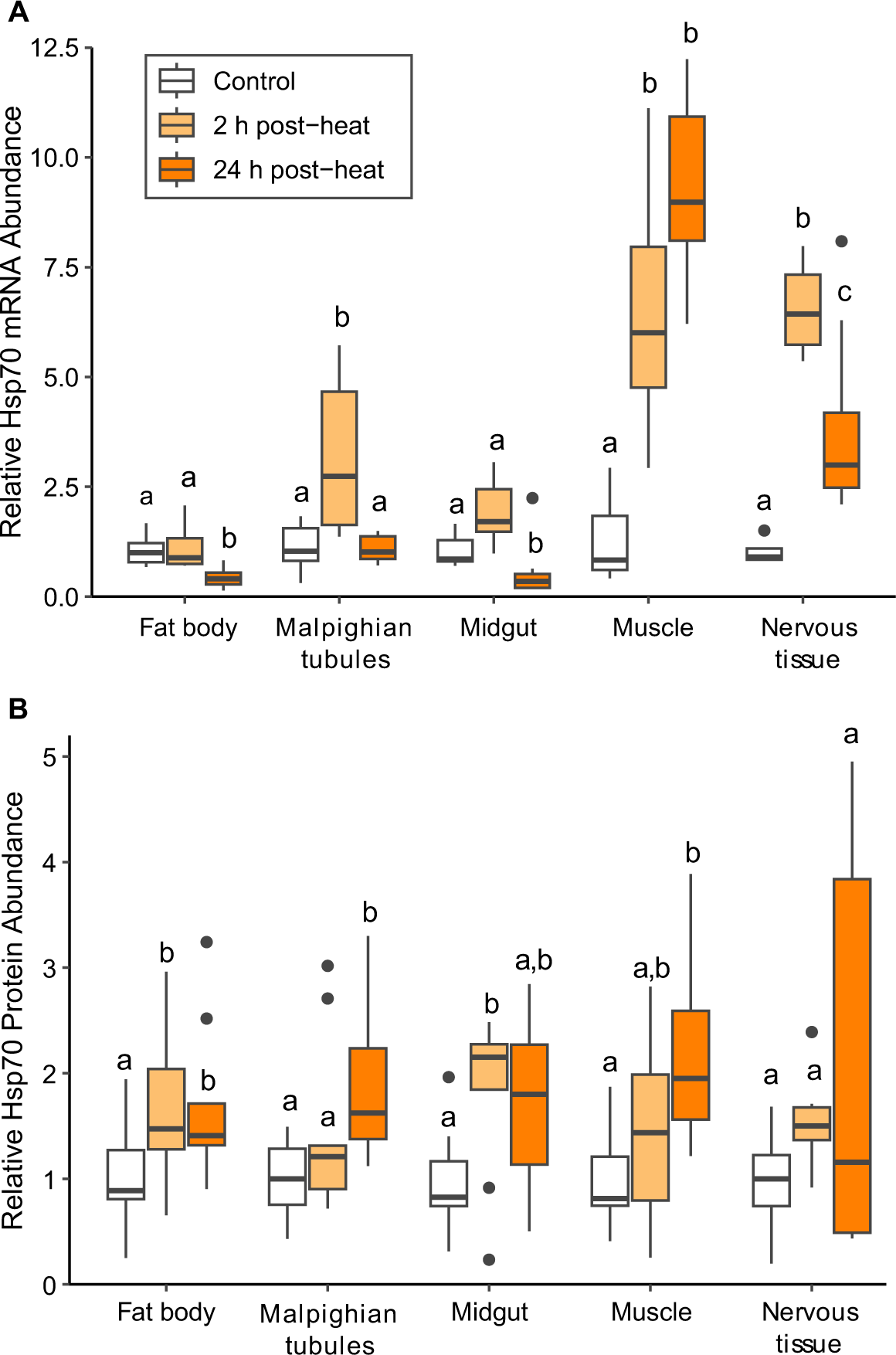
Relative HSP70 (A) mRNA and (B) protein abundance in five tissues of *Gryllus veletis* that were not acclimated and not heat shocked (white; controls) or heat shocked (40°C for 2 h) and recovered for 2 h (light orange) or 24 h (dark orange). Black dots represent outliers. Within a tissue, different letters indicate a significant difference in HSP70 mRNA abundance (ANOVA with Tukey’s test, *P* < 0.05) or protein abundance (Kruskal-Wallis with Dunn’s test, *P* < 0.05) relative to controls.

**Table 3.**
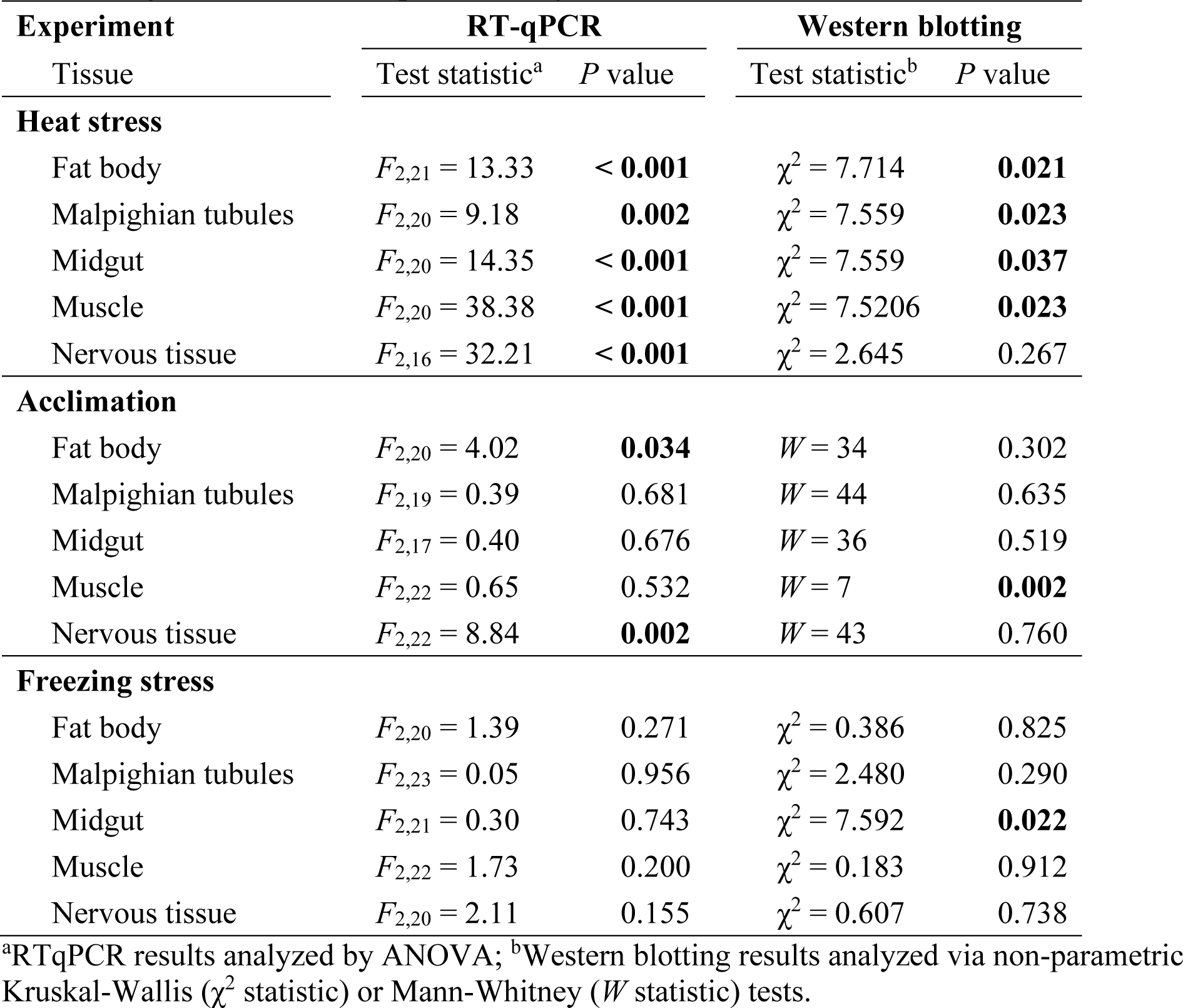
Statistical results comparing relative abundance of HSP70 mRNA (RT-qPCR) and protein (Western blotting) in three experiments (Figure 1) that tested the effect of different thermal regimes on HSP70 expression. Significant *P*-values are bolded.

| Experiment | RT-qPCR |  | Western blotting |  |
| --- | --- | --- | --- | --- |
| Tissue | Test statistic<sup>a</sup> | *P* value | Test statistic<sup>b</sup> | *P* value |
| **Heat stress** |  |  |  |  |
| Fat body | $F_{2,21} = 13.33$ | **< 0.001** | $\chi^2 = 7.714$ | **0.021** |
| Malpighian tubules | $F_{2,20} = 9.18$ | **0.002** | $\chi^2 = 7.559$ | **0.023** |
| Midgut | $F_{2,20} = 14.35$ | **< 0.001** | $\chi^2 = 7.559$ | **0.037** |
| Muscle | $F_{2,20} = 38.38$ | **< 0.001** | $\chi^2 = 7.5206$ | **0.023** |
| Nervous tissue | $F_{2,16} = 32.21$ | **< 0.001** | $\chi^2 = 2.645$ | 0.267 |
| **Acclimation** |  |  |  |  |
| Fat body | $F_{2,20} = 4.02$ | **0.034** | $W = 34$ | 0.302 |
| Malpighian tubules | $F_{2,19} = 0.39$ | 0.681 | $W = 44$ | 0.635 |
| Midgut | $F_{2,17} = 0.40$ | 0.676 | $W = 36$ | 0.519 |
| Muscle | $F_{2,22} = 0.65$ | 0.532 | $W = 7$ | **0.002** |
| Nervous tissue | $F_{2,22} = 8.84$ | **0.002** | $W = 43$ | 0.760 |
| **Freezing stress** |  |  |  |  |
| Fat body | $F_{2,20} = 1.39$ | 0.271 | $\chi^2 = 0.386$ | 0.825 |
| Malpighian tubules | $F_{2,23} = 0.05$ | 0.956 | $\chi^2 = 2.480$ | 0.290 |
| Midgut | $F_{2,21} = 0.30$ | 0.743 | $\chi^2 = 7.592$ | **0.022** |
| Muscle | $F_{2,22} = 1.73$ | 0.200 | $\chi^2 = 0.183$ | 0.912 |
| Nervous tissue | $F_{2,20} = 2.11$ | 0.155 | $\chi^2 = 0.607$ | 0.738 |

<sup>a</sup>RTqPCR results analyzed by ANOVA; <sup>b</sup>Western blotting results analyzed via non-parametric Kruskal-Wallis ( $\chi^2$ statistic) or Mann-Whitney ( $W$ statistic) tests.

### HSP70 was upregulated in fat body, femur muscle, and nervous tissue during acclimation

The relative abundance of HSP70 mRNA or protein abundance significantly increased during fall-like acclimation in fat body, femur muscle, and nervous tissue (Figure 6, Table 3), but not as substantially as in the heat shock experiment. Target HSP70 mRNA abundance was approximately 2-to 2.5-fold higher in fat body and nervous tissue after 6 weeks of acclimation compared to unacclimated crickets (Figure 6A). At the protein level, only femur muscle tissue had a significant increase in relative HSP70 abundance (Figure 6B).

**Figure 6.**
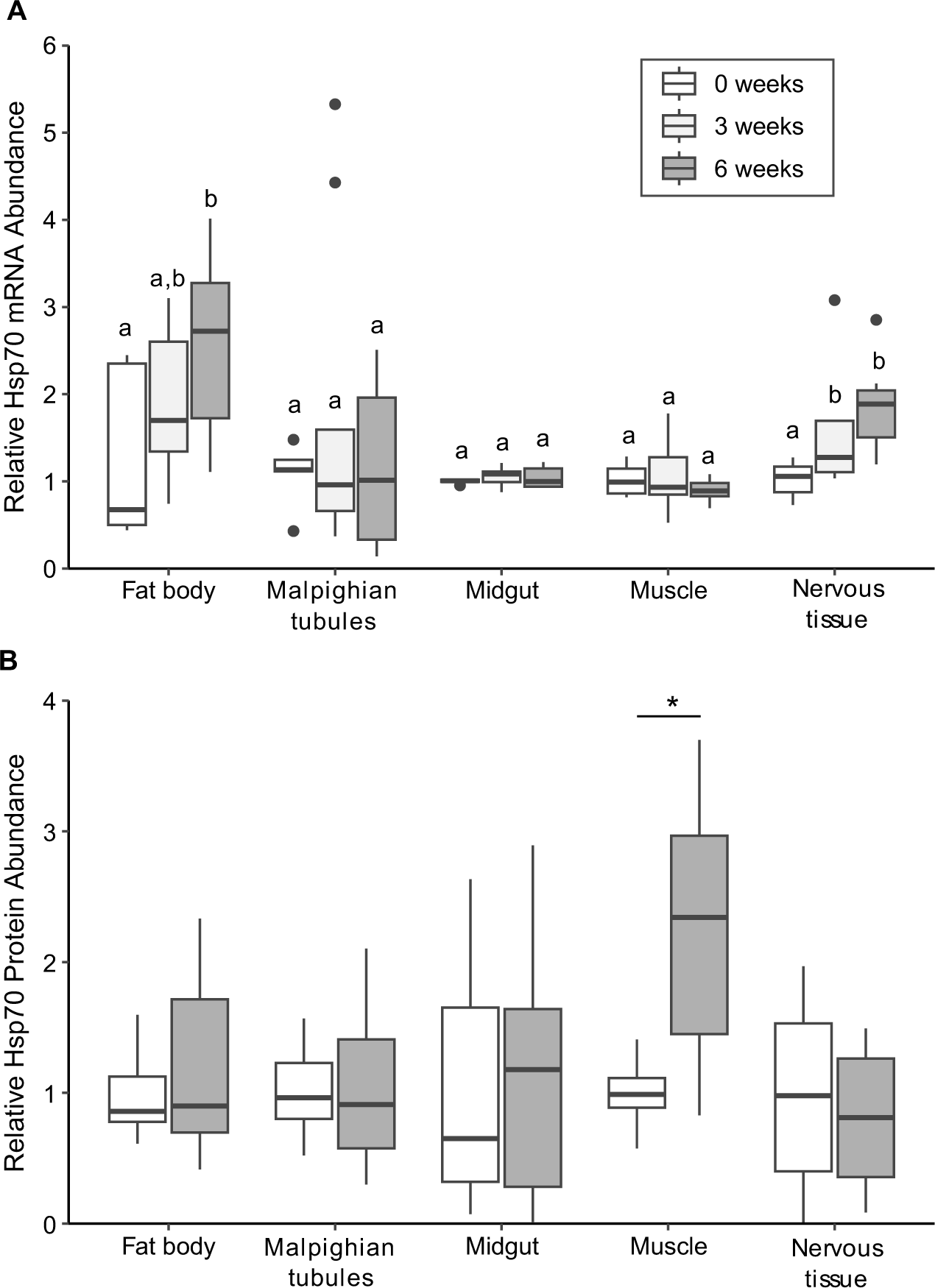
Relative HSP70 (A) mRNA and (B) protein abundance in five tissues of *Gryllus veletis* after 0 (white; controls), 3 (light grey), and 6 (dark grey) weeks of fall-like acclimation. Black dots represent outliers. Within a tissue, different letters or asterisks indicate a significant difference in HSP70 mRNA abundance (ANOVA with Tukey’s test, *P* < 0.05) or protein abundance (Mann-Whitney test, *P* < 0.05) relative to controls. 3 week acclimation samples were not available for Western blotting, so are not represented in panel **(B)**.

### Freezing did not induce upregulation of HSP70

HSP70 mRNA and protein expression was not significantly different after the freezing stress in any tissue except midgut, which showed an unexpected decrease in relative abundance of HSP70 protein (Figure 7, Table 3). Although we expected high survival after the freeze treatment, only 30 of the 54 crickets (56%) of frozen and thawed crickets were alive at the time of dissection (similar proportions 2 h and 24 h post-freeze), potentially caused by the crickets recovering at room temperature (c. 20°C) post-freeze, rather than the moderate temperature of 15°C that has been used previously (Toxopeus et al., 2019c). If there was upregulation of HSP70 in surviving crickets, the signal was too weak to cause an increase in the average HSP70 abundance, so we conclude there was not a strong effect of freezing on HSP70 expression.

**Figure 7.**
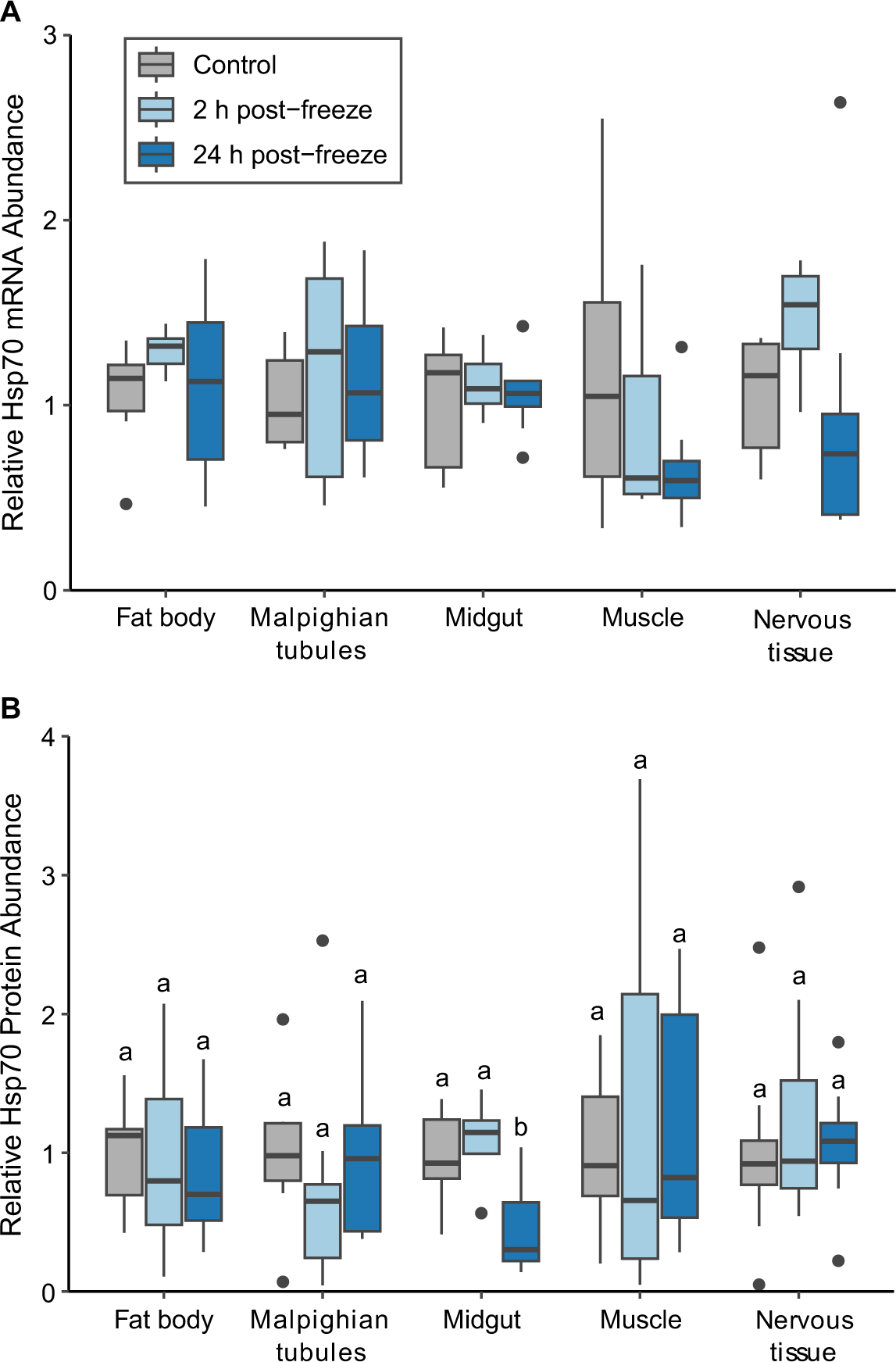
Relative HSP70 (A) mRNA and (B) protein abundance in five tissues of *Gryllus veletis* that were acclimated and never frozen (dark gray; controls) or frozen (-8°C for 1.5 h) and recovered for 2 h (light blue) or 24 h (dark blue). Black dots represent outliers. Within a tissue, there was no significant difference in HSP70 mRNA abundance (ANOVA *P* > 0.05); and different letters indicate significant differences in HSP70 protein abundance (Kruskal-Wallis with Dunn’s test, *P* < 0.05) relative to controls.

## Discussion

While freeze tolerance has been studied in *G. veletis* (Smith et al., 2021; Toxopeus et al., 2019a, 2019b, 2019c), this paper is the first to describe survival following heat stress, expanding our perspective on the stress tolerance of this laboratory model. Although heat stress could induce HSP70 expression at the mRNA and/or protein level in all tissues, we saw mild upregulation of HSP70 during the fall-like acclimation that induces freeze tolerance in only three of the five tissues, and no increase in HSP70 expression following freezing stress in any tissues. These results provide relatively weak support for our hypothesis that HSP70 is important for freeze tolerance, despite its importance in the response to heat stress. We propose two possible interpretations of these results, which we discuss in more detail below: 1) the HSP70 or other chaperones accumulated during fall-like acclimation were sufficient to mitigate any damage to proteins due to freezing and thawing; 2) freezing stress didn’t cause protein damage, and therefore HSP70 was not required for post-freeze recovery.

First, we must establish that our target HSP70s in *G. veletis* were stress-inducible, which was done via our heat stress experiment. The survivable heat shock (40°C for 2 h) we used was sufficient to stimulate HSP70 upregulation at the mRNA and protein level, suggesting that the heat shock caused sufficient protein denaturation or damage to induce a response by molecular chaperones (Storey and Storey, 2023; Ullah et al., 2024). Although the HSP70 encoded by Gvel_34771 had higher similarity to HSC70 than HSP70 sequences (see Supplementary Information), the mRNA for this putative HSP70 was indeed heat-inducible in three of the five tissues we studied – nervous tissue, femur muscle, and Malpighian tubules. Others have also shown that HSC70 mRNA can be upregulated by heat stress (Fang et al., 2021; Koštál and Tollarová-Borovanská, 2009; Li et al., 2021), and our results are consistent with other findings of tissue-specific expression of HSP isoforms (Fang et al., 2021; Wang et al., 2019; Yocum, 2001). Based on an overall increase in relative abundance of HSP70 protein in the other tissues we studied – fat body and midgut – we suggest that HSP70 isoforms other than Gvel_34771 are also heat-inducible and exhibit tissue-specific expression patterns. Future work could examine mRNA abundance of other potential HSP70 isoforms in the *G. veletis* transcriptome (Toxopeus et al., 2019a) to understand the heat shock response more fully in this species.

HSP70 or other chaperone upregulation at the mRNA or protein level in *G. veletis* during fall-like acclimation may have been sufficient to protect against or repair protein damage caused during freezing, thus minimizing the need for HSP70 upregulation following freezing stress. Our results are similar to the whole-body transcriptomics in freeze-tolerant *C. costata*, which also upregulate HSP70 during acclimation but not following freezing (Des Marteaux et al., 2019).

The upregulation of HSP70 mRNA that we saw in fat body during acclimation was consistent with previous transcriptomic results in *G. veletis* (Toxopeus et al., 2019a), and paralleled by changes in the nervous tissue. However, at the protein level we only saw a notable increase in HSP70 in femur muscle during acclimation, suggesting the crickets prioritize protection of this tissue and likely use a different HSP70 isoform than that encoded by Gvel_34771. While HSP70 upregulation may be an important preparatory mechanism for freeze tolerance in some tissues, the lack of upregulation in other tissues indicates this mechanism is not universal, and highlights the important of tissue-specific work (e.g., Des Marteaux et al., 2017). Investigation into other inducible chaperones (e.g., Cai et al., 2017; Des Marteaux et al., 2019) that are cold-responsive could be an interesting avenue for future research.

It is possible that freezing was insufficiently stressful to cause protein denaturation or damage, making it unnecessary for *G. veletis* to upregulate HSP70 after freezing. The freeze treatment we used (-8°C for 1.5 h) can be considered relatively mild for our study species; acclimated *G. veletis* can have a high survival rate (>90%) if frozen for up to 48 h at this temperature (Toxopeus et al., 2019c). In addition, the freeze treatment we used included gradual decreases and increases to and from the target temperature (respectively), which can minimize the challenges associated with thermal stress (Bahar et al., 2013; Hemmati et al., 2014; but see Sørensen et al., 2013). This is in contrast to our heat stress experiment, in which crickets were exposed to high temperatures without any gradual temperature ramps. Grgac et al. (2022) have shown that even lethal freeze treatments do not impair soluble enzyme (protein) function in a range of insect species, suggesting that freezing does not damage these proteins. More stressful low temperature exposures such as repeated freeze-thaw events (e.g., Gill et al., 2023) may be required to induce upregulation of HSP70 in *G. veletis*. To test whether freezing causes protein damage in *G. veletis*, additional studies that examine enzyme activity post-freeze (as in Grgac et al., 2022) or other markers of protein damage and aggregation such as ubiquitination (as in Gill et al., 2023) would be required.

## Conclusion

Although HSP70 is important for responding to a wide range of stressors, our work suggests that this chaperone may be less critical in the preparation for, and response to, moderate freezing stress in a freeze-tolerant insect. While we could detect HSP70 upregulation following heat stress at 40°C, only a subset of tissues upregulated HSP70 as *G. veletis* underwent acclimation to become freeze-tolerant, and no tissue exhibited HSP70 upregulation after a short freezing treatment at -8°C. This first detailed study of HSP70 in *G. veletis* prompts further questions about the intensity and frequency of freezing stress that can cause cellular protein damage, and the extent to which mechanisms that protect proteins are important for surviving freezing.

## Data and Code Availability

All data and code associated with this manuscript are available via the GitHub repository: https://github.com/jtoxopeus/cricket-hsp70

## Acknowledgements

The authors would like to thank P. Brown, A. Gough. L. McIntyre, M. Moore, and the StFX Animal Care Facility for help with rearing and/or processing crickets for our experiments.

## Author contributions

**VEA**: conceptualization, methodology, investigation (RT-qPCR), analysis, writing – original draft, writing – review and editing. **MLvO**: methodology, investigation (Western blotting), analysis, writing – original draft, writing – review and editing. **JT**: conceptualization, data curation, funding acquisition, investigation (sequence analysis), supervision, writing – original draft, writing – review and editing.

