## Supplementary Information for "HSP70 is upregulated after heat but not freezing stress in the freeze-tolerant cricket *Gryllus veletis*"

**Table of Contents**

Explanatory text: Target selection for RT-qPCR and protein structure analysis.............................2

Table S1: Putative HSP70 transcripts from *Gryllus veletis* ...........................................................3

Figure S1: Alignment of putative protein sequence of *Gryllus veletis* HSP70 transcript Gvel_43771_c0_g1 with other insect protein sequences.............................................4-6

Figure S2: Partial alignment of putative *Gryllus veletis* HSP70 Gvel_43771_c0_g1 against other orthopteran HSP and HSC nucleotide sequences ............................................7-12

Figure S3: Alignment of putative *Gryllus veletis* HSP70 transcripts to each other..................13-21

Figure S4: Partial alignment of putative *Gryllus veletis* HSP70 transcripts Gvel_19946_c0_g1, Gvel_16103_c0_g1 and Gvel_83898_c0_g1_i1 against other orthopteran sequences........................................................................22-29

**Target selection for RT-qPCR and protein structure analysis**

Six putative transcripts from the *Gryllus veletis* transcriptome (Toxopeus et al. 2019a) were therefore potential targets for RT-qPCR of HSP70 in our study due to their high similarity to other orthopteran HSP70 sequences (Table S1). Of these six transcripts, Gvel_34771 (NCBI accession GGSD01051771.1) was sufficiently long (2,470 bp; Table S1) to include the full ORF (open reading frame) for a HSP70 sequence. The other five putative transcripts were all shorter (< 750 bp; Table S1), and therefore likely represented fragments of a full mRNA transcript and were not as suitable for further analysis.

We also suggest that Gvel_34771 represents a different isoform of HSP70 than the other five putative transcript fragments. The longest ORF from Gvel_34771 encoded a putative protein that aligned well with HSC70-4 protein sequences from other insects (Figure S1), similar to the nucleotide sequence (Table S1, Figure S2). For its protein sequence, Gvel_34771 showed high (99%) identity to *Gryllus bimaculatus* HSC70-4 (Figure S1). For its nucleotide sequence, Gvel_34771 had high identity to *Schistocerca gregaria* HSC70-4 (80%), and lower identity with *Schistocerca americana* Hsp70-B2 (58%; Table S1, Figure S2). Conversely, while the other five putative HSP transcript fragments could align with Gvel_34771 (albeit with < 70% identity; Figure S3), these five transcript fragments had the highest identity (>80%) to an inducible HSP70 isoform: HSP70-B2 from *Schistocerca* species (Table S1, Figure S4). The transcript fragments also showed some alignment to each other with high identity than to Gvel_34771. There was overlap between Gvel_19946 and Gvel_16103 (89% identity), Gvel_78482_c1 and Gvel_78482_c2 (97% identity), Gvel_78482_c1/2 and Gvel_83898 (83% identity; Figure S3).

**Table S1**. Putative HSP70 transcripts from *Gryllus veletis* (*G. vel*) were identified by BLASTing three known locust HSP70 sequences (*Schistocerca americana* XM_047140202.1, XM_047139358.1, and MT498301.1) against the Transcriptome Shotgun Assembly (TSA) database of *G. vel* on NCBI. Each putative HSP70 transcript was then BLASTed against the Standard nucleotide database on NCBI. *G. vel* transcripts > 500 bp and BLAST hits with high query coverage and % identity were potential targets for RT-qPCR.

| **Putative *G. vel* HSP70 transcript (NCBI Accession; description[a])** | **Length (bp)** | **Top BLAST hit (NCBI Accession; *Species*[b], Gene[c])** | **% Query Cover** | **% Identity** | **E-value** | **Potential target?** |
|---|---|---|---|---|---|---|
| GGSD01051771.1; Gvel_34771_c0_g1_i1 | 2,470 | XM_049987033.1; *Schistocera gregaria* HSC70-4 | 80 | 80 | 0.0 | Yes |
| GGSD01027946.1; Gvel_19946_c0_g1_i1 | 734 | XM_050106196.1; *Schistocerca serialis* HSP70-B2 | 94 | 84 | 0.0 | Yes |
| GGSD01022798.1; Gvel_16103_c0_g1_i1 | 504 | XM_050106196.1; *Schistocerca serialis* HSP70-B2 | 98 | 83 | 2e-123 | Yes |
| GGSD01130040.1; Gvel_83898_c0_g1_i1 | 746 | XM_049928178.1; *Schistocerca cancellata* HSP70-B2 | 89 | 84 | 3e-172 | Yes |
| GGSD01119730.1; Gvel_78482_c1_g1_i1 | 289 | XM_050106196.1; *Schistocerca serialis* HSP70-B2 | 97 | 92 | 2e-105 | Maybe, but short |
| GGSD01119729.1; Gvel_78482_c2_g1_i1 | 306 | XM_049955302.1; *Schistocerca nitens* HSP70-B2 | 99 | 84 | 5e-72 | Maybe, but short |
| GGSD01094007.1; Gvel_61233_c0_g1_i1 | 375 | XM_022369462.2; *Drosophila obscura* HSP70 Ba | 96 | 73 | 5e-23 | No – short, not Orthoptera |
| GGSD01026446.1; Gvel_18794_c0_g1_i1 | 262 | XM_034393654.1; *Thrips palmi* HSP70 Ab | 96 | 88 | 5e-76 | No – short; not Orthoptera |
| GGSD01089422.1; Gvel_57949_c0_g1_i1 | 236 | XM_020620091.1; *Monopterus albus* HSC71 | 27 | 94 | 5e-16 | No - short, low query cover |
| GGSD01089421.1; Gvel_57949_c0_g2_i1 | 237 | XM_020620091.1; *Monopterus albus* HSC71 | 27 | 94 | 5e-16 | No - short, low query cover |
| GGSD01089420.1; Gvel_57949_c0_g3_i1 | 238 | XM_020620091.1; *Monopterus albus* HSC71 | 27 | 94 | 5e-16 | No - short, low query cover |
| GGSD01135028.1; Gvel_87722_c0_g1_i1 | 5,450 | XM_046539502.1; *Ischnura elegans* HSP70-4 | 27 | 75 | 6e-177 | No - low query cover |
| GGSD01135027.1; Gvel_87722_c0_g1_i2 | 5,474 | XM_046539502.1; *Ischnura elegans* HSP70-4 | 27 | 75 | 6e-177 | No - low query cover |

[a]Description is the transcript ID in Toxopeus et al. (2019a); [b]*Schistocerca* is the only orthopteran genus (same order as *Gryllus*) in this column; [c]HSP = heat shock protein, HSC = heat shock cognate.

**Figure S1 (1/2)**

```
                         cov     pid   1 [         .         .         .         .         .         . 60
1 Gvel_34771_ORF     100.0% 100.0%      MALKAPAVGIDLGTTYSCVGVFQHGKVEIIANDQGNRTTPSYVAFTETERLIGDAAKNQV
2 G.bimac_Hsc70-4     99.5%  99.5%      MALKAPAVGIDLGTTYSCVGVFQHGKVEIIANDQGNRTTPSYVAFTETERLIGDAAKNQV
3 S.gregaria_HSP70    99.1%  92.7%      MAVKAPAVGIDLGTTYSCVGVFQHGKVEIIANDQGNRTTPSYVAFTDTERLIGDAAKNQV
4 D.mel_Hsc70-4       98.6%  86.3%      -MSKAPAVGIDLGTTYSCVGVFQHGKVEIIANDQGNRTTPSYVAFTDTERLIGDAAKNQV
5 D.mel_Hsp70Ab       97.1%  74.0%      ----MPAIGIDLGTTYSCVGVYQHGKVEIIANDQGNRTTPSYVAFTDSERLIGDPAKNQV

                         cov     pid  61           .         .         .         .         .         . 120
1 Gvel_34771_ORF     100.0% 100.0%      AMNPNNTIFDAKRLIGRRFEDQTVQADMKHWPFNVISDSGKPKIQVQYKGESKTFYPEEI
2 G.bimac_Hsc70-4     99.5%  99.5%      AMNPNNTIFDAKRLIGRRFEDQTVQADMKHWPFNVISDSGKPKIQVQYKGESKTFYPEEI
3 S.gregaria_HSP70    99.1%  92.7%      AMNPSNTIFDAKRLIGRRFDDQAVQSDMKHWPFKVISDAGKPKIQVQYKGETKTFFPEEV
4 D.mel_Hsc70-4       98.6%  86.3%      AMNPTQTIFDAKRLIGRKFDDAAVQSDMKHWPFEVVSADGKPKIEVTYKDEKKTFFPEEI
5 D.mel_Hsp70Ab       97.1%  74.0%      AMNPRNTVFDAKRLIGRKYDDPKIAEDMKHWPFKVVSDGGKPKIGVEYKGESKRFAPEEI

                         cov     pid 121           .         .         .         .         .         . 180
1 Gvel_34771_ORF     100.0% 100.0%      SSMVLTKMKETAEAYLGKNVTNAVITVPAYFNDSQRQATKDAGAIAGLNVLRIINEPTAA
2 G.bimac_Hsc70-4     99.5%  99.5%      SSMVLTKMKETAEAYLGKNVTNAVITVPAYFNDSQRQATKDAGAIAGLNVLRIINEPTAA
3 S.gregaria_HSP70    99.1%  92.7%      SSMVLTKMKETAEAYLGKNVSNAVITVPAYFNDSQRQATKDAGAIAGLNVLRIINEPTAA
4 D.mel_Hsc70-4       98.6%  86.3%      SSMVLTKMKETAEAYLGKTVTNAVITVPAYFNDSQRQATKDAGTIAGLNVLRIINEPTAA
5 D.mel_Hsp70Ab       97.1%  74.0%      SSMVLTKMKETAEAYLGESITDAVITVPAYFNDSQRQATKDAGHIAGLNVLRIINEPTAA

                         cov     pid 181           .         .         .         .         .         . 240
1 Gvel_34771_ORF     100.0% 100.0%      AIAYGLDKKVSNGHGERNVLIFDLGGGTFDVSILTIEDG-IFEVKSTAGDTHLGGEDFDN
2 G.bimac_Hsc70-4     99.5%  99.5%      AIAYGLDKK---GHGERNVLIFDLGGGTFDVSILTIEDG-IFEVKSTAGDTHLGGEDFDN
3 S.gregaria_HSP70    99.1%  92.7%      AIAYGLDKK---GHGERNVLIFDLGGGTFDVSILTIEDG-IFEVKATAGDTHLGGEDFDN
4 D.mel_Hsc70-4       98.6%  86.3%      AIAYGLDKK---AVGERNVLIFDLGGGTFDVSILSIDDG-IFEVKSTAGDTHLGGEDFDN
5 D.mel_Hsp70Ab       97.1%  74.0%      ALAYGLDKN---LKGERNVLIFDLGGGTFDVSILTIDEGSLFEVRSTAGDTHLGGEDFDN

                         cov     pid 241           .         .         .         .         .           300
1 Gvel_34771_ORF     100.0% 100.0%      RMVNHFVQEFKRKYKKDLATNKRALRRLRTACERAKRTLSSSTQASIEIDSLFEGIDFYT
2 G.bimac_Hsc70-4     99.5%  99.5%      RMVNHFVQEFKRKYKKDLATNKRALRRLRTACERAKRTLSSSTQASIEIDSLFEGIDFYT
3 S.gregaria_HSP70    99.1%  92.7%      RMVNHFVQEFKRKYKKDLTTNKRALRRLRTACERAKRTLSSSTQASIEIDSLYEGIDFYT
4 D.mel_Hsc70-4       98.6%  86.3%      RLVTHFVQEFKRKHKKDLTTNKRALRRLRTACERAKRTLSSSTQASIEIDSLFEGTDFYT
5 D.mel_Hsp70Ab       97.1%  74.0%      RLVTHLADEFKRKYKKDLRSNPRALRRLRTAAERAKRTLSSSTEATIEIDALFEGQDFYT

                         cov     pid 301           .         .         .         .         .         . 360
1 Gvel_34771_ORF     100.0% 100.0%      SITRARFEELNADLFRSTMEPVEKSLRDAKIDKAQIHDIVLVGGSTRIPKVQKLLQDFFN
2 G.bimac_Hsc70-4     99.5%  99.5%      SITRARFEELNADLFRSTMEPVEKSLRDAKIDKAQIHDIVLVGGSTRIPKVQKLLQDFFN
3 S.gregaria_HSP70    99.1%  92.7%      SITRARFEELNADLFRSTMEPVEKALRDAKMDKAQIHDIVLVGGSTRIPKVQKLLQDFFN
4 D.mel_Hsc70-4       98.6%  86.3%      SITRARFEELNADLFRSTMDPVEKALRDAKLDKSVIHDIVLVGGSTRIPKVQRLLQDLFN
5 D.mel_Hsp70Ab       97.1%  74.0%      KVSRARFEELCADLFRNTLQPVEKALNDAKMDKGQIHDIVLVGGSTRIPKVQSLLQDFFH
```

**Figure S1 (2/2)**

```
                          cov    pid 361      .         .         .         .         . 420
1 Gvel_34771_ORF      100.0% 100.0%     GKELNKSINPDEAVAYGAAVQAAILAGDKSEEVQDLLLLDVTPLSLGIETAGGVMTALIK
2 G.bimac_Hsc70-4      99.5%  99.5%     GKELNKSINPDEAVAYGAAVQAAILAGDKSEEVQDLLLLDVTPLSLGIETAGGVMTALIK
3 S.gregaria_HSP70     99.1%  92.7%     GKELNKSINPDEAVAYGAAVQAAILAGDKSEEVQDLLLLDVTPLSLGIETAGGVMTTLIK
4 D.mel_Hsc70-4        98.6%  86.3%     GKELNKSINPDEAVAYGAAVQAAILHGDKSQEVQDLLLLDVTPLSLGIETAGGVMSVLIK
5 D.mel_Hsp70Ab        97.1%  74.0%     GKNLNLSINPDEAVAYGAAVQAAILSGDQSGKIQDVLLVDVAPLSLGIETAGGVMTKLIE

                          cov    pid 421      .         .         .         .         . 480
1 Gvel_34771_ORF      100.0% 100.0%     RNTTIPTKQTQTFTTYSDNQPGVLIQVYEGERAMTKDNNLLGKFELTGIPPAPRGVPQIE
2 G.bimac_Hsc70-4      99.5%  99.5%     RNTTIPTKQTQTFTTYSDNQPGVLIQVYEGERAMTKDNNLLGKFELTGIPPAPRGVPQIE
3 S.gregaria_HSP70     99.1%  92.7%     RNTTIPTKQTQTFTTYSDNQPGVLIQVYEGERAMTKDNNLLGKFELTGIPPAPRGVPQIE
4 D.mel_Hsc70-4        98.6%  86.3%     RNTTIPTKQTQTFTTYSDNQPGVLIQVYEGERAMTKDNNLLGKFELSGIPPAPRGVPQIE
5 D.mel_Hsp70Ab        97.1%  74.0%     RNCRIPCKQTKTFSTYADNQPGVSIQVYEGERAMTKDNNALGTFDLSGIPPAPRGVPQIE

                          cov    pid 481      .         .         .         .         . 540
1 Gvel_34771_ORF      100.0% 100.0%     VTFDIDANGILNVTAIEKSTGKENKITITNDKGRLSKEEIERMVNDAEKYRAEDDKQKQT
2 G.bimac_Hsc70-4      99.5%  99.5%     VTFDIDANGILNVTAIEKSTGKENKITITNDKGRLSKEEIERMVNDAEKYRAEDDKQKQT
3 S.gregaria_HSP70     99.1%  92.7%     VTFDIDANGILNVTAVEKSTGKENKITITNDKGRLSKEEIERMVNEAERYRAEDEKQKAT
4 D.mel_Hsc70-4        98.6%  86.3%     VTFDIDANGILNVTALERSTNKENKITITNDKGRLSKEDIERMVNEAEKYRNEDEKQKET
5 D.mel_Hsp70Ab        97.1%  74.0%     VTFDLDANGILNVSAKEMSTGKAKNITIKNDKGRLSQAEIDRMVNEAEKYADEDEKHRQR

                          cov    pid 541      .         .         .         .         6 600
1 Gvel_34771_ORF      100.0% 100.0%     IAAKNSLESYCFNMKSTVEDEKLKDKISDADKTSILDKCNEVIRWLDSNQLAEKEEFEAQ
2 G.bimac_Hsc70-4      99.5%  99.5%     IAAKNSLESYCFNMKSTVEDEKLKDKISDADKTSILDKCNEVIRWLDSNQLAEKEEFEAQ
3 S.gregaria_HSP70     99.1%  92.7%     IAAKNGLESYCFNMKSTVEDEKLKDKISDSDKQTILDKCNEVIRWLDANQLAEKEEFEEK
4 D.mel_Hsc70-4        98.6%  86.3%     IAAKNGLESYCFNMKATLDEDNLKTKISDSDRTTILDKCNETIKWLDANQLADKEEYEHR
5 D.mel_Hsp70Ab        97.1%  74.0%     ITSRNALESYVFNVKQAVEQA-PAGKLDEADKNSVLDKCNDTIRWLDSNTTAEKEEFDHK

                          cov    pid 601      .         .         .         .         . 660
1 Gvel_34771_ORF      100.0% 100.0%     QKELEALCNPIITKLYQGAGGAPGGMP-GFPGGFPGGGAAPGPGGAAGAGAGAGPTIEEV
2 G.bimac_Hsc70-4      99.5%  99.5%     QKELEALCNPIITKLYQGAGGAPGGMP-GFPGGFPGGGAAPGPGGAAGAGAGAGPTIEEV
3 S.gregaria_HSP70     99.1%  92.7%     QKELEQICNPIITKLYQGAGGAPGGMPGGFPGGFPGAGGAAAGGAG---AGGAGPTIEEV
4 D.mel_Hsc70-4        98.6%  86.3%     QKELEGVCNPIITKLYQGAGFPPGGMPGG-PGGMPGAAGAAG----AAGAGGAGPTIEEV
5 D.mel_Hsp70Ab        97.1%  74.0%     LEELTRHCSPIMTKMHQQGAGAGAGGPGANCGQQ-----------AGGFGGYSGPTVEEV

                          cov    pid 661 ] 661
1 Gvel_34771_ORF      100.0% 100.0%     D
2 G.bimac_Hsc70-4      99.5%  99.5%     D
3 S.gregaria_HSP70     99.1%  92.7%     D
4 D.mel_Hsc70-4        98.6%  86.3%     D
5 D.mel_Hsp70Ab        97.1%  74.0%     D
```

**Figure S1 (above)**. Alignment of putative protein sequence of *Gryllus veletis* HSP70 transcript GGSD01051771.1 (Gvel_43771_c0_g1) with other HSP70 protein sequences from NCBI (GLG99747.1; *Gryllus bimaculatus* HSC70-4) and UniProt: A0A8E5JSY6 (*Schistocerca gregaria* HSP70), P11147 (*Drosophila melanogaster* HSC70-4), P02825 (*Drosophila melanogaster* HSP70Ab). Alignment performed with Clustal Omega and visualized with MView. Amino acids that match the *G. veletis* sequence are colour-coded. Percent coverage (cov) and percent identity (pid) for each sequence relative to the *G. veletis* sequence are reported, showing highest identity with HSC70-4 sequences.

**Figure S2 (1/5)**

```
Gvel_34771_c0_g1         CGCTATTATCTAAAGAAAATGGCTCTCAAAGCGCCTGCGGTTGGTATTGATCTTGGAACT    294
S.gregaria.Hsc70         ATACGCTAACGAAAGAAAATGGCTGTTAAAGCACCTGCAGTGGGAATTGATCTAGGTACC    1177
S.americana.Hsp70A1      TGACATAGCTAACAGAAGAAAATGCCGAAGATTCCAGCAGTGGGGATCGATTTGGGAACC    244
S.americana.HSP70B2      CAGCAGCCATCAGCAGCGACCATGGGCAAGGCAACCGCCGTCGGCATAGACCTGGGCACC    88
                                    *      *        **     * ** ** ** ** **  * ** **

Gvel_34771_c0_g1         ACCTATTCCTGTGTTGGTGTTTTTCAACACGGGAAGGTAGAAATCATTGCCAATGACCAG    354
S.gregaria.Hsc70         ACCTACTCCTGTGTTGGAGTGTTCCAGCATGGGAAAGTGGAAATCATCGCCAATGATCAA    1237
S.americana.Hsp70A1      ACGTACTCGTGCGTGGGAGTGTGGCAACAAGGCAAGGTGGAAATAATCGCCAACGATCAA    304
S.americana.HSP70B2      ACCTACTCGTGCGTGGGCGTGTGGCAGCACGGCAAGGTCGAGATCGTCGCCAACGAGCAG    148
                         ** ** ** ** ** ** ** *  ** ** ** ** ** ** **  * ***** ** **

Gvel_34771_c0_g1         GGAAACAGAACCACGCCCAGCTATGTTGCATTTACTGAGACTGAGCGCCTTATTGGCGAT    414
S.gregaria.Hsc70         GGGAATCGTACAACACCTAGCTACGTCGCATTTACGGACACAGAGCGACTCATTGGCGAT    1297
S.americana.Hsp70A1      GGCAACAGGACGACACCAAGCTACGTCGCCTTTTGCGACTCGGAGCGGCTGATCGGCGAT    364
S.americana.HSP70B2      GGCAACCGCACCACGCCCAGCTACGTCGCCTTCACCGACACCGAGCGACTCATCGGCGAT    208
                         ** **  * ** ** ** ***** ** ** **    **  * ***** ** ** ******

Gvel_34771_c0_g1         GCTGCCAAAAATCAAGTGGCCATGAACCCCAATAACACAATTTTTGATGCCAAACGTCTT    474
S.gregaria.Hsc70         GCTGCTAAAAATCAGGTGGCCATGAACCCAAGTAACACTATTTTTGATGCAAAGCGTCTT    1357
S.americana.Hsp70A1      GCTGCCAAGAACCAGGTTGCGATGAACCCGCAGAATACCATCTTCGACGCCAAACGACTC    424
S.americana.HSP70B2      GCCGCCAAAAGCCAGGTGGCCATGAACCCGAAGAACACGGTTTTCGACGCGAAGCGATTG    268
                         ** ** ** *  ** ** ** ********    ** **  * ** ** ** ** **  *

Gvel_34771_c0_g1         ATTGGGAGAAGATTTGAAGATCAAACTGTTCAAGCCGACATGAAACATTGGCCATTTAAT    534
S.gregaria.Hsc70         ATTGGGCGCCGTTTCGACGACCAGGCTGTACAAAGCGATATGAAGCATTGGCCTTTCAAA    1417
S.americana.Hsp70A1      ATCGGCCGCAAGTTTGACGACCCAAAGGTGCAGGGAGACATGAAGCATTGGCCATTCAAA    484
S.americana.HSP70B2      ATTGGAAGACGCTTTGACGATCCCAAGATACAAGACGACATGAAGCACTGGCCGTTCACG    328
                         ** **  *    ** ** ** *      * **    ** ***** ** ***** ** *

Gvel_34771_c0_g1         GTAATTAGCGACAGTGGAAAACCAAAGATCCAAGTTCAGTACAAAGGAGAATCTAAAACT    594
S.gregaria.Hsc70         GTCATAAGTGATGCTGGCAAACCAAAGATTCAGGTTCAGTACAAAGGCGAAACGAAGACT    1477
S.americana.Hsp70A1      GTGATTAACGACTGTAGCAAGCCTAAGATACAGGTCCAGTTCAAAGGCACCACCAAAACT    544
S.americana.HSP70B2      GTCGTTTCCGACGGCGACAAACCCAAAATTCAGGTGGAGTACAAGGGCGAGACGAAGAGG    388
                         **  *    **       ** ** ** ** ** **  *** *** **     * ** *

Gvel_34771_c0_g1         TTTTACCCTGAAGAAATTAGTTCCATGGTCCTCACAAAGATGAAGGAAACAGCAGAGGCT    654
S.gregaria.Hsc70         TTCTTCCCTGAGGAGGTTAGCTCAATGGTCCTGACAAAAATGAAGGAAACTGCAGAGGCG    1537
S.americana.Hsp70A1      TTCGCTCCAGAAGAGATAAGCGCTATGGTGCTTGTGAAAATGAAGGAAACAGCAGAGGCA    604
S.americana.HSP70B2      TTCGCGCCCGAGGAGATCAGCTCGATGGTGCTGAGCAAGATGCGCGAGATCGCGAGCACG    448
                         **    ** ** **  * **  * ***** **    ** ***   ** *  **     *
```

**Figure S2 (2/5)**

```
Gvel_34771_c0_g1           TATTTGGGCAAGAATGTTACAAATGCTGTGATTACCGTTCCTGCTTATTTCAATGATTCC    714
S.gregaria.Hsc70           TACCTTGGTAAAAATGTCAGCAACGCTGTGATTACGGTTCCTGCCTACTTCAATGATTCG    1597
S.americana.Hsp70A1        TTCTTGGGTGGGCAGGTTCGAGAGGCAGTTATTACAGTGCCCGCCTACTTCAACGACTCA    664
S.americana.HSP70B2        TACCTGGGCGGCGACGTGCGCGACGCGGTGATCACGGTGCCGGCCTACTTCAACGACGCG    508
                           *   * **     * **     * ** ** ** ** ** ** ** ** ***** **  *

Gvel_34771_c0_g1           CAAAGACAAGCCACAAAGGATGCTGGTGCCATTGCTGGTTTGAATGTTCTTCGCATAATC    774
S.gregaria.Hsc70           CAGAGGCAAGCCACCAAGGATGCGGGAGCCATTGCTGGACTCAATGTGTTGCGAATTATT    1657
S.americana.Hsp70A1        CAGCGGCAAGCTACCAAAGATGCAGGCGCCATAGCAGGCCTGAAGGTGCTGCGGATTATC    724
S.americana.HSP70B2        CAGCGCAGCGCCACCAAGGACGCGGGCGCCATCGCCGGCCTCAACGTGCTGCGCATCATC    568
                           **  *    ** ** ** ** ** ** ***** ** **  * ** **  * ** ** **

Gvel_34771_c0_g1           AATGAGCCTACAGCTGCAGCCATTGCTTATGGCCTCGACAAAAAGGTGAGTAATGGTCAT    834
S.gregaria.Hsc70           AACGAGCCAACAGCGGCTGCAATTGCCTATGGTCTCGATAAGAAGGTAA---GTGGTCAT    1714
S.americana.Hsp70A1        AACGAGCCAACTGCAGCTGCACTCGCATACGGCCTCGACAAAAACCTGAA---------A    775
S.americana.HSP70B2        AACGAGCCCACCGCCGCCGCGCTCGCCTACGGGCTCGACAAGAACCTGCA---------G    619
                           ** ***** ** ** ** **  * ** ** ** ***** ** **  *

Gvel_34771_c0_g1           GGTGAGAGAAATGTTCTCATTTTTGACCTGGGTGGCGGCACCTTCGATGTATCAATTCTG    894
S.gregaria.Hsc70           GGTGAAAGAAACGTGCTTATCTTTGACTTGGGTGGTGGTACATTTGATGTGTCAATTCTG    1774
S.americana.Hsp70A1        GGTGAGCGCAATGTGCTCATATTTGACCTTGGTGGTGGCACCTTTGACGTGTCCATCCTG    835
S.americana.HSP70B2        GGCGAGAAGAACGTGCTCATCTTCGACCTCGGCGGAGGCACTTTTGACGTGTCGGTGCTG    679
                           ** **    ** ** ** ** ** *** * ** ** ** ** ** ** ** **  * ***

Gvel_34771_c0_g1           ACCATTGAA---GATGGAATCTTTGAAGTGAAATCTACAGCTGGAGACACTCATCTGGGT    951
S.gregaria.Hsc70           ACAATTGAA---GATGGTATTTTTGAAGTGAAAGCCACAGCAGGAGACACTCATTTAGGA    1831
S.americana.Hsp70A1        AGCATTTCGGAGGGTTCACTGTTTGAAGTGAAAGCAACAGCTGGAGACACACACCTGGGA    895
S.americana.HSP70B2        GCCATCTCGGAAGGGTCGCTGTTCGAGGTGAAGTCGACGGCGGGTGATACGCACCTGGGC    739
                              **       *      * ** ** *****  * ** ** ** ** ** **  * **

Gvel_34771_c0_g1           GGTGAAGACTTCGATAACAGAATGGTGAACCACTTTGTTCAAGAATTCAAGAGGAAATAC    1011
S.gregaria.Hsc70           GGTGAAGACTTTGATAACCGCATGGTTAACCATTTTGTGCAAGAATTTAAGAGAAAGTAC    1891
S.americana.Hsp70A1        GGCGAGGACTTTGACAGTCGGCTTGTGAATCATTTGGCTGATGAGTTCAAACGTAAATTC    955
S.americana.HSP70B2        GGCGAGGACTTCGACAACCGGCTGGTGCAGCACCTGGCGGAAGAGTTCCAGCGCAAGCAC    799
                           ** ** ***** ** *   *  * **  *  **  * *   * ** **  *  * **   *

Gvel_34771_c0_g1           AAGAAAGATCTGGCAACCAACAAGAGAGCTCTTCGTCGCCTACGTACTGCCTGTGAAAGA    1071
S.gregaria.Hsc70           AAAAAAGACCTTACTACCAACAAAAGAGCACTGCGTAGGTTGAGAACTGCCTGTGAAAGA    1951
S.americana.Hsp70A1        CACAAGGACGTACGTTCCAATCCACGTGCTTTGCGTCGACTGCGTACGGCAGCAGAACGG    1015
S.americana.HSP70B2        CGCAAGGACATGCGCGCCAACGCGCGCGCGCTGCGCCGCCTGCGCACCGCCGCCGAGCGC    859
                              ** **  *     ****     * **  * **  *  *  * ** **    **  *
```

**Figure S2 (3/5)**

```
Gvel_34771_c0_g1          GCAAAGCGTACTTTATCTTCGTCAACTCAAGCCAGTATTGAAATTGATTCTCTCTTTGAG   1131
S.gregaria.Hsc70          GCAAAACGCACCCTATCTTCGTCAACTCAAGCCAGTATTGAAATAGATTCTCTCTATGAG   2011
S.americana.Hsp70A1       GCCAAGCGTACGTTGTCTTCCAGCACGGAGGCCAGTATTGAGATTGATGCACTATTAGAT   1075
S.americana.HSP70B2       GCCAAGCGGACGCTCTCGTCCAGCACCGAGGCCAGCCTCGAGATAGACGCCCTGCACGAC   919
                          ** ** ** **  * ** **    **  * *****  * ** ** **  * **    **

Gvel_34771_c0_g1          GGAATAGATTTCTACACTTCTATCACTAGGGCTCGCTTTGAAGAATTGAATGCTGACCTC   1191
S.gregaria.Hsc70          GGTATTGACTTCTATACATCTATAACAAGAGCAAGGTTTGAAGAGCTCAATGCTGACTTG   2071
S.americana.Hsp70A1       GGCATTGACTTCTACACAAAAGTTTCACGAGCACGGTTTGAGGAACTTTGTGCTGATCTG   1135
S.americana.HSP70B2       GGCATCGACTTCTACGCCAAGGTGACGCGCGCGGTTCGAGGAGCTGTGCATGGACCTC   979
                          ** ** ** *****  *     *  *  * **  * ** ** **  *       **  *

Gvel_34771_c0_g1          TTCCGCAGCACCATGGAACCTGTTGAGAAGTCTTTGCGTGACGCAAAGATCGACAAGGCT   1251
S.gregaria.Hsc70          TTCAGGTCAACTATGGAACCTGTGGAGAAAGCACTCCGTGATGCGAAGATGGACAAAGCT   2131
S.americana.Hsp70A1       TTCCGCTCAACACTACAACCTGTCGAGAAAGCACTGGCCGATGCCAAGATGGACAAAGCG   1195
S.americana.HSP70B2       TTCCGGCAGACGCTGGCGCCGGTGGAGCGCGCCCTGGGCGACGCCAAGCTGGACAAGGCG   1039
                          *** *    **  *    ** ** ***    *  *    ** ** *** * ***** **

Gvel_34771_c0_g1          CAAATTCACGACATTGTTCTTGTTGGTGGTTCTACTCGCATTCCTAAGGTTCAAAAGCTC   1311
S.gregaria.Hsc70          CAGATTCATGACATCGTGTTGGTTGGTGGGTCAACTCGTATTCCAAAAGTACAAAAGTTG   2191
S.americana.Hsp70A1       TCAATCCACGATGTCGTGTTGGTGGGTGGTTCAACACGCATCCCTAAGATTCAGGCTCTG   1255
S.americana.HSP70B2       TCCGTGCACGACGTCGTGCTGGTCGGCGGCTCCACGCGCATCCCCAAGATACAGAAGATG   1099
                              * ** **  * **  * ** ** ** ** ** ** ** ** **  * **     *

Gvel_34771_c0_g1          CTACAAGATTTCTTCAATGGCAAAGAATTGAACAAATCCATCAACCCTGATGAAGCTGTT   1371
S.gregaria.Hsc70          CTTCAAGACTTCTTCAATGGCAAAGAACTGAACAAATCAATTAACCCAGATGAGGCAGTA   2251
S.americana.Hsp70A1       CTGCAGAACTACTTTGCTGGTAAGCGGCTGAATTTGTCCATCAACCCGGATGAGGCAGTG   1315
S.americana.HSP70B2       CTGCAGGACTTCTTCTGCGGCAAGACGCTGAACCTGTCCATCAACCCGGACGAGGCGGTG   1159
                          ** **  * * ***    ** **     ****    ** ** ***** ** ** ** **

Gvel_34771_c0_g1          GCTTATGGTGCTGCTGTGCAGGCTGCAATTTTGGCTGGTGATAAGTCTGAGGAGGTTCAA   1431
S.gregaria.Hsc70          GCATATGGAGCAGCTGTACAAGCAGCAATATTGGCAGGTGACAAGTCTGAAGAAGTGCAG   2311
S.americana.Hsp70A1       GCATATGGTGCAGCTGTACAAGCAGCCATTCTCAGTGGTGATACCAGCTCTGCAATTCAG   1375
S.americana.HSP70B2       GCGTACGGCGCGGCGGTGCAGGCGGCCATCCTGAGCGGCGACACGAGCTCGCAGATCCAG   1219
                          ** ** ** ** ** ** ** ** ** **  *    ** ** *            * **

Gvel_34771_c0_g1          GATCTCCTTTTGCTTGATGTTACACCATTGTCTCTGGGTATTGAAACTGCTGGAGGAGTC   1491
S.gregaria.Hsc70          GACTTGCTACTGCTTGATGTTACACCACTATCTCTTGGTATAGAGACTGCAGGTGGTGTG   2371
S.americana.Hsp70A1       GATGTGTTACTGGTAGATGTAGCACCACTATCACTCGGCATTGAGACAGCAGGTGGTGTC   1435
S.americana.HSP70B2       GACGTGCTGCTGGTCGACGTGGCGCCGCTCTCGCTCGGCATCGAGACCGCGGGCGGCGTC   1279
                          **  *  *  ** * ** **  * **  * ** ** ** ** ** ** ** ** ** **
```

**Figure S2 (4/5)**

```
Gvel_34771_c0_g1          ATGACAGCCCTCATCAAGCGAAACACAACCATTCCAACCAAGCAGACCCAAACTTTCACC   1551
S.gregaria.Hsc70          ATGACGACTCTTATCAAACGAAATACCACAATCCCAACAAAGCAAACCCAGACATTCACA   2431
S.americana.Hsp70A1       ATGACTAAGCTGGTAGAGCGCAATGCGCGTATCCCCTGCAAGCAAAAACAGACATTCACC   1495
S.americana.HSP70B2       ATGACCAAGATCGTCGAGCGCAACGCGCGCATCCCCTGCAAGCAGAAGCAGACCTTCACC   1339
                          *****     *  *  * ** **  *    ** **    ***** *  ** ** *****

Gvel_34771_c0_g1          ACATATTCTGATAACCAACCTGGTGTACTCATCCAGGTTTACGAGGGTGAAAGAGCCATG   1611
S.gregaria.Hsc70          ACATATTCGGACAACCAACCTGGTGTGCTCATTCAGGTGTATGAAGGTGAGCGTGCCATG   2491
S.americana.Hsp70A1       ACATACTCAGACAACCAGCCAGCAGTTACAATTCAGGTGTTTGAAGGAGAGCGTGCAATG   1555
S.americana.HSP70B2       ACCTACTCCGACAACCAGCCCGCCGTCACCATTCAGGTGTTCGAGGGCGAGCGCGCCATG   1399
                          ** ** ** ** ***** ** *  **    ** ***** *  ** ** **  * ** ***

Gvel_34771_c0_g1          ACCAAGGACAACAACCTTCTTGGAAAGTTTGAACTGACTGGCATTCCCCCAGCTCCACGT   1671
S.gregaria.Hsc70          ACAAAGGACAACAATCTTCTGGGTAAATTCGAATTGACAGGAATTCCACCTGCCCCAAGA   2551
S.americana.Hsp70A1       ACAAAGGACAACAACTTGCTGGGCACATTTGACCTCACTGGCATTCCACCAGCACCACGT   1615
S.americana.HSP70B2       ACCAAGGACAACAACCTGCTGGGCACGTTCAACCTGACGGGCATCCCGCCGGCGCCGCGG   1459
                          ** ***********  * ** ** *  **  *  * ** ** ** ** ** ** **  *

Gvel_34771_c0_g1          GGTGTCCCACAAATTGAAGTTACTTTTGATATTGATGCTAATGGTATTTTGAATGTAACT   1731
S.gregaria.Hsc70          GGTGTGCCTCAAATTGAAGTCACTTTTGACATTGATGCTAATGGTATCTTGAATGTAACT   2611
S.americana.Hsp70A1       GGTGTGCCTCAAGTGGAAGTTACATTTGACTTGAATGCAGATGGCATTCTAACTGTGTCC   1675
S.americana.HSP70B2       GGAGTACCCAAGATCGAGGTGACCTTCGACCTGGACGCGAACGGCATCCTCAACGTGTCG   1519
                          ** ** **  *  * ** ** ** ** **  * ** ** ** ** **  * *  **  *

Gvel_34771_c0_g1          GCTATTGAGAAATCTACTGGAAAAGAAAACAAGATCACAATCACCAATGACAAGGGTCGT   1791
S.gregaria.Hsc70          GCTGTGGAGAAATCAACTGGCAAGGAGAACAAAATCACAATTACAAATGACAAGGGCCGT   2671
S.americana.Hsp70A1       GCCCAAGAGAAGAGCACAGGAAGATCTCGTAACATCACCATCCGCAATGACAAGGGTCGG   1735
S.americana.HSP70B2       GCGACGGAAAGCGGCTCGGGTCGCAGCGAGCGCATCACCATCCAGAACGACAAGGGCCGC   1579
                          **    ** *      * **             ***** **    ** ******** **

Gvel_34771_c0_g1          CTTAGCAAGGAAGAAATAGAGCGTATGGTTAATGATGCTGAAAAATACCGTGCTGAAGAT   1851
S.gregaria.Hsc70          TTGAGCAAGGAAGAAATCGAGCGCATGGTCAATGAAGCTGAACGTTACCGTGCTGAAGAC   2731
S.americana.Hsp70A1       CTTTCCAAGGAGGAAATCGATCGGATGGTAGCTGATGCAGAGCGCTTCAAGGAGGAAGAC   1795
S.americana.HSP70B2       CTCTCCAAGGCCGAGATCGAGCGCATGCTGGCCGACGCAGAGCGCTTCCGCGCCGAGGAC   1639
                           *   *****  ** ** ** ** *** *    ** ** **    * *   *  ** **

Gvel_34771_c0_g1          GACAAACAGAAACAGACCATTGCTGCTAAGAATTCCTTGGAATCCTACTGTTTCAACATG   1911
S.gregaria.Hsc70          GAGAAGCAGAAGGCAACTATTGCAGCGAAAAATGGACTCGAGTCTTACTGCTTCAACATG   2791
S.americana.Hsp70A1       CAGCGGCAACGTGAGAGAGTAGAGGCACGGCATCGACTGGAGGGCTATGCCCTGTCTGTG   1855
S.americana.HSP70B2       GAGCGGCAGCGCGCGCGGGTCGAGGCCCGCAACCGCCTGGAAGCGTACGCGCTCTCGCTC   1699
                           *    **           * *  **     *     * **    **     *     *
```

**Figure S2 (5/5)**

```
Gvel_34771_c0_g1         AAGAGCACTGTTGAAGACGAGAAGCTGAAGGATAAGATCTCTGATGCTGATAAGACTTCA   1971
S.gregaria.Hsc70         AAGTCTACTGTGGAAGACGAAAAGTTGAAAGACAAGATCTCCGACTCAGACAAGCAAACT   2851
S.americana.Hsp70A1      AAGCAGGCACTGTCGGACGC------AGGCAGCAGGCTATCTGAGTCGGACAAGACAGCA   1909
S.americana.HSP70B2      AAGCAGGCCGCCGAGGACGC------GGGCAGCAAGCTGAGCGACGCGGACAAGGCGACG   1753
                         ***    *       ****              * * *    **  * ** ***    *

Gvel_34771_c0_g1         ATTTTGGACAAGTGCAATGAGGTCATTCGCTGGCTGGATTCTAACCAACTTGCTGAAAAA   2031
S.gregaria.Hsc70         ATCTTGGACAAGTGCAATGAAGTTATCCGTTGGCTTGATGCCAACCAGCTGGCTGAGAAG   2911
S.americana.Hsp70A1      GCGATGACACGTTGTGAGGAGGAATTACAGTGGCTAGACAACAATGCCCAGGCGAGCTGT   1969
S.americana.HSP70B2      GTGCGGGACCAGAGCGCGGAGGCGCTCAAGTGGCTGGACAGCAACTCGCTGGCGGAGCAG   1813
                              *       *    ** *   *    ***** **    **    *  **

Gvel_34771_c0_g1         GAAGAATTCGAGGCCCAACAGAAGGAACTTGAAGCGCTCTGCAACCCAATAATCACCAAG   2091
S.gregaria.Hsc70         GAAGAATTTGAGGAGAAGCAGAAGGAACTCGAGCAGATCTGCAATCCTATCATCACCAAA   2971
S.americana.Hsp70A1      GGTGAGTGTGAGCGTCGGCTACAGGAGCTGCAGCGGGTATGCACACCTATAATGACTAAG   2029
S.americana.HSP70B2      GAGGAGTTCGAGGACCGCTACAAGCAGCTGTCGGCCGCCTGCTCGCCCATCATGGCCAAG   1873
                         *  ** *  ***          ** * **          ***   ** ** **  * **

Gvel_34771_c0_g1         TTGTACCAGGGTGCTGGTGGAGCTCCTGGTGGCATGCCTGGATTTCCTGGTGGT---TTC   2148
S.gregaria.Hsc70         TTGTACCAGGGTGCAGGAGGAGCTCCTGGAGGAATGCCTGGTGGTTTCCCTGGTGGATTC   3031
S.americana.Hsp70A1      CTGCACCAGGGCACTAAGGATCCCAGCAGTAGCTGTGGGCAACAAGCAGGTACAGGAGCA   2089
S.americana.HSP70B2      CTGCACCAGGCGGGCGCCGGCGGCGGCGCGGGCGCAGGC------------ACGGGCTCC   1921
                          ** ******        *            *

Gvel_34771_c0_g1         CCTGGTGGCGGTGCCGCTCCTGGACCTGGTGGT---GCTGCTGGTGCTGGAGCCGGTGCT   2205
S.gregaria.Hsc70         CCAGGTGCTGGTGGAGCTGC---------TGCT---GGTGGTGCAGGTGCCGGTGGTGCT   3079
S.americana.Hsp70A1      GCACACACCGGTGGACCAACTATTGAGGAGGTTGACTAACTGA-----------------   2132
S.americana.HSP70B2      GGGGCCGCCAAGGGCCCCACCGTCGAGGAGGTCGACTGAGGCGGCGGCGAGGCCGCTCCG   1981
                                    *   *  *          *

Gvel_34771_c0_g1         GGACCTACTATTGAAGAAGTTGATTAAACAATTCATCACTTGCAACCATTT----TAGTT   2261
S.gregaria.Hsc70         GGGCCAACTATTGAAGAGGTTGACTAAATGTGAAAGTGCTAAAGGTTAATCTGTCCAGTT   3139
S.americana.Hsp70A1      ----------------AC------------------------------------------   2134
S.americana.HSP70B2      TGAAGAACTCTACTGGACTGTGGCCACACCATTCTAAC-------ACACTCCTTACACTA   2034
                                         *

Gvel_34771_c0_g1         CATGGCTAAAGTTTCATTCCAGAATCAGTATTATTGCAGACACAACTTAAAGTCAATTTT   2321
S.gregaria.Hsc70         CAGT-GCAGTTTTCAGTTCAAGCTTCAAGATTTTTGTTAACTTGA---------------   3183
S.americana.Hsp70A1      ---TGTTAATATAAAAACGG--------TTTTACGTGTCACTGTGTGTGATATATAAAAT   2183
S.americana.HSP70B2      GCCTGCCACTCTCGTTTGGGAATTACTTCGTTGTGCGTCACACACTATGCTCCAAATTTT   2094
                                *   *                  **       **
```

**Figure S2 (above)**. Partial alignment of putative *Gryllus veletis* HSP70 transcript GGSD01051771.1 (Gvel_43771_c0_g1) against other orthopteran HSP and HSC nucleotide sequences from NCBI: XM_049987033.1 (*Schistocerca gregaria* Hsc70), XM_047139358.1 (*S. americana* Hsp70-A1), and XM_047140202.1 (*S. americana* Hsp70-B2). Produced using Clustal Omega. Asterisks indicate the same nucleotide in all four sequences.

**Figure S3 (1/9)**

```
Gvel_34771_c0_g1      GAAATATGTAATTTTAAGTACCTGCTTTGGTCGATTGTCGCGCGAACATTCGGCGTGACG   60
Gvel_19946_c0_g1      ------------------------------------------------------------   0
Gvel_16103_c0_g1      ------------------------------------------------------------   0
Gvel_83898_c0_g1      ------------------------------------------------------------   0
Gvel_78482_c1_g1      ------------------------------------------------------------   0
Gvel_78482_c2_g1      ------------------------------------------------------------   0

Gvel_34771_c0_g1      TTTTGCTTTCCCATTGGCCGGATGGATAGAAGTTTCTAGAAGCTTGTGGTCGTCATCGAC   120
Gvel_19946_c0_g1      ------------------------------------------------------------   0
Gvel_16103_c0_g1      ------------------------------------------------------------   0
Gvel_83898_c0_g1      ------------------------------------------------------------   0
Gvel_78482_c1_g1      ------------------------------------------------------------   0
Gvel_78482_c2_g1      ------------------------------------------------------------   0

Gvel_34771_c0_g1      GTTATAAAAGAGAGGTCGGCCGCCGACGCCCCAGCAGTACAAGTTAGTATAGCGGAGTGT   180
Gvel_19946_c0_g1      ------------------------------------------------------------   0
Gvel_16103_c0_g1      ------------------------------------------------------------   0
Gvel_83898_c0_g1      ------------------------------------------------------------   0
Gvel_78482_c1_g1      ------------------------------------------------------------   0
Gvel_78482_c2_g1      ------------------------------------------------------------   0

Gvel_34771_c0_g1      GAAGTTAAGTTAAGTTGTGTGAATTCGTCTTCGAGTTTTGTCTGTAATTCAAAGCGCTAT   240
Gvel_19946_c0_g1      ------------------GCGGAGGACG-------CGGCGCCCGAGAGCGACATTTGAAC   35
Gvel_16103_c0_g1      ------------------------------------------------------------   0
Gvel_83898_c0_g1      ------------------------------------------------------------   0
Gvel_78482_c1_g1      ------------------------------------------------------------   0
Gvel_78482_c2_g1      ------------------------------------------------------------   0
                                                             . . .  .   . .     .

Gvel_34771_c0_g1      TATCTAAAGAAAATGGCTCTCAAAGCGCCTGCGGTTGGTATTGATCTTGGAACTACCTAT   300
Gvel_19946_c0_g1      AGCATTTCAGCCACCATGGGGAAAGTGCCTGCAGTGGGAATTGATCTCGGCACGACCTAT   95
Gvel_16103_c0_g1      ------------------------------------------------------------   0
Gvel_83898_c0_g1      ------------------------------------------------------------   0
Gvel_78482_c1_g1      ------------------------------------------------------------   0
Gvel_78482_c2_g1      ------------------------------------------------------------   0
                          .       .        .... ...... .. .. ........ .. .. ......
```

**Figure S3 (2/9)**

```
Gvel_34771_c0_g1    TCCTGTGTTGGTGTTTTTCAACACGGGAAGGTAGAAATCATTGCCAATGACCAGGGAAAC  360
Gvel_19946_c0_g1    TCGTGCGTCGGGGTCTGGCAGCATGGTAAAGTAGAGATAATCGCCAATGACCAAGGAAAT  155
Gvel_16103_c0_g1    ------------------------------------------------------------  0
Gvel_83898_c0_g1    ------------------------------------------------------------  0
Gvel_78482_c1_g1    ------------------------------------------------------------  0
Gvel_78482_c2_g1    ------------------------------------------------------------  0
                    .. .. .. .. .. .  .. .. .. .. ..... .. .. ........... ..... 

Gvel_34771_c0_g1    AGAACCACGCCCAGCTATGTTGCATTTACTGAGACTGAGCGCCTTATTGGCGATGCTGCC  420
Gvel_19946_c0_g1    CGCACAACACCCAGTTATGTCGCTTTCACGGACACTGAACGGCTTATCGGCGATGCTGCC  215
Gvel_16103_c0_g1    ------------------------------------------------------------  0
Gvel_83898_c0_g1    ------------------------------------------------------------  0
Gvel_78482_c1_g1    ------------------------------------------------------------  0
Gvel_78482_c2_g1    ------------------------------------------------------------  0
                       .. .. ..... ..... .. .. .. .. ..... .. ..... ............

Gvel_34771_c0_g1    AAAAATCAAGTGGCCATGAACCCCAATAACACAATTTTTGATGCCAAACGTCTTATTGGG  480
Gvel_19946_c0_g1    AAAAGTCAGGTGGCGATGAACCCAAAGAACACCGTTTTCGACGCCAAACGCTTAATAGGA  275
Gvel_16103_c0_g1    ------------------------------------------------------------  0
Gvel_83898_c0_g1    ------------------------------------------------------------  0
Gvel_78482_c1_g1    ------------------------------------------------------------  0
Gvel_78482_c2_g1    ------------------------------------------------------------  0
                    .... ... ..... ........ .. .....  .... .. ........  . .. .. 

Gvel_34771_c0_g1    AGAAGATTTGAAGATCAAACTGTTCAAGCCGACATGAAACATTGGCCATTTAATGTAATT  540
Gvel_19946_c0_g1    AGGCGCTTCGATGATGCGAAGATTCAAGACGACATGAAGCACTGGCCGTTTACTGTCGTC  335
Gvel_16103_c0_g1    ------------------------------------------------------------  0
Gvel_83898_c0_g1    ------------------------------------------------------------  0
Gvel_78482_c1_g1    ------------------------------------------------------------  0
Gvel_78482_c2_g1    ------------------------------------------------------------  0
                    ..  . .. .. ...   .   ...... ......... .. ..... .... ...  . 

Gvel_34771_c0_g1    AGCGACAGTGGAAAACCAAAGATCCAAGTTCAGTACAAAGGAGAATCTAAAACTTTTTAC  600
Gvel_19946_c0_g1    AACGATGGCGGCAAGCCCAAGATCCAGGTGGAGTACAAGAGCGAGGTGAAGCGCTTCGCG  395
Gvel_16103_c0_g1    ------------------------------------------------------------  0
Gvel_83898_c0_g1    ------------------------------------------------------------  0
Gvel_78482_c1_g1    ------------------------------------------------------------  0
Gvel_78482_c2_g1    ------------------------------------------------------------  0
                    . ...  . .. .. .. ........ ..  .......  . ..    ..     .. 
```

**Figure S3 (3/9)**

```
Gvel_34771_c0_g1      CCTGAAGAAATTAGTTCCATGGTCCTCACAAAGATGAAGGAAACAGCAGAGGCTTATTTG  660
Gvel_19946_c0_g1      CCGGAGGAGGTGAGCTCCATGGTGCTGACCAAGATGCGCGAGATCGCCGAGACGTACCTG  455
Gvel_16103_c0_g1      ------------------------------------------------------------  0
Gvel_83898_c0_g1      ------------------------------------------------------------  0
Gvel_78482_c1_g1      ------------------------------------------------------------  0
Gvel_78482_c2_g1      ------------------------------------------------------------  0
                      .. .. ..  . .. ........ .. .. ......   .. .  .. ... . ..  ..

Gvel_34771_c0_g1      GGCAAGAATGTTACAAATGCTGTGATTACCGTTCCTGCTTATTTCAATGATTCCCAAAGA  720
Gvel_19946_c0_g1      GGCGGGAAGGTGACGGACGCGGTGATCACGGTGCCGGCCTACTTCAACGACTCGCAGCGG  515
Gvel_16103_c0_g1      ------------------------------------------------------------  0
Gvel_83898_c0_g1      ------------------------------------------------------------  0
Gvel_78482_c1_g1      ------------------------------------------------------------  0
Gvel_78482_c2_g1      ------------------------------------------------------------  0
                      ...  ... .. ..  . .. ..... .. .. .. .. .. ..... .. .. ..  .

Gvel_34771_c0_g1      CAAGCCACAAAGGATGCTGGTGCCATTGCTGGTTTGAATGTTCTTCGCATAATCAATGAG  780
Gvel_19946_c0_g1      CAGGCCACCAAGGACGCCGGCGCCATCGCGGGCCTCAACGTGCTGCGGATCATCAACGAG  575
Gvel_16103_c0_g1      ------------------------------------------------------------  0
Gvel_83898_c0_g1      ------------------------------------------------------------  0
Gvel_78482_c1_g1      ------------------------------------------------------------  0
Gvel_78482_c2_g1      ------------------------------------------------------------  0
                      .. ..... ..... .. .. ..... .. ..  . .. .. .. .. .. ..... ...

Gvel_34771_c0_g1      CCTACAGCTGCAGCCATTGCTTATGGCCTCGACAAAAAGGTGAGTAATGGTCATGGTGAG  840
Gvel_19946_c0_g1      CCGACGGCCGCCGCGCTGGCCTACGGCCTCGACAAGAACCTCAA---------GGGCGAG  626
Gvel_16103_c0_g1      ------------------------------------------------------------  0
Gvel_83898_c0_g1      ------------------------------------------------------------  0
Gvel_78482_c1_g1      ------------------------------------------------------------  0
Gvel_78482_c2_g1      ------------------------------------------------------------  0


Gvel_34771_c0_g1      AGAAATGTTCTCATTTTTGACCTGGGTGGCGGCACCTTCGATGTATCAATTCTGACCATT  900
Gvel_19946_c0_g1      AAGAACGTGCTCATCTTCGACCTCGGCGGCGGCACCTTCGACGTCTCCATCCTCTCCATC  686
Gvel_16103_c0_g1      ------------------------------------------------------------  0
Gvel_83898_c0_g1      ------------------------------------------------------------  0
Gvel_78482_c1_g1      ------------------------------------------------------------  0
Gvel_78482_c2_g1      ------------------------------------------------------------  0
                      .  .. .. ..... .. ..... .. .............. .. .. .. ..  ....
```

**Figure S3 (4/9)**

```
Gvel_34771_c0_g1      GAAGATGGAATCTT---TGAAGTGAAATCTACAGCTGGAGACACTCATCTGGGTGGTGAA  957
Gvel_19946_c0_g1      GACGAGGGATCGCTCTTCGAGGTGAAGTCGACGGCCGGCGACACGCAC------------  734
Gvel_16103_c0_g1      -CCGAGGGCTCGCTCTTCGAGGTGCGGGCGACGGCCGGCGACACGCACCTGGGCGGCGAG  59
Gvel_83898_c0_g1      ------------------------------------------------------------  0
Gvel_78482_c1_g1      ------------------------------------------------------------  0
Gvel_78482_c2_g1      ------------------------------------------------------------  0
                      ...::.::....:....::.:::....:.::.::.::.:::::.::......  .. .. 

Gvel_34771_c0_g1      GACTTCGATAACAGAATGGTGAACCACTTTGTTCAAGAATTCAAGAGGAAATACAAGAAA  1017
Gvel_19946_c0_g1      ------------------------------------------------------------  734
Gvel_16103_c0_g1      GACTTCGACACGCGCCTCGTCGCCCACCTCGCCGACGAGTTCCGGCGCAAGCACGGCAAG  119
Gvel_83898_c0_g1      ------------------------------------------------------------  0
Gvel_78482_c1_g1      ------------------------------------------------------------  0
Gvel_78482_c2_g1      ------------------------------------------------------------  0
                      ........ .      . ..   .... . .   . .. ...  . . ..  ..   .. 

Gvel_34771_c0_g1      GATCTGGCAACCAACAAGAGAGCTCTTCGTCGCCTACGTACTGCCTGTGAAAGAGCAAAG  1077
Gvel_19946_c0_g1      ------------------------------------------------------------  734
Gvel_16103_c0_g1      GACGTGCGCCGCCACGCGCGCGCCCTGCGCCGCCTGCGCACCGCCGCCGAGCGCGCCAAG  179
Gvel_83898_c0_g1      ------------------------------------------------------------  0
Gvel_78482_c1_g1      ------------------------------------------------------------  0
Gvel_78482_c2_g1      ------------------------------------------------------------  0
                      ..  ..     .  .  . . .. .. .. ..... .. .. ...   ..  . .. ...

Gvel_34771_c0_g1      CGTACTTTATCTTCGTCAACTCAAGCCAGTATTGAAATTGATTCTCTCTTTGAGGGAATA  1137
Gvel_19946_c0_g1      ------------------------------------------------------------  734
Gvel_16103_c0_g1      CGCACCCTCTCCTCCAGCACCGAGGCCGGCGTCGAGGTGGACGCCCTGCTCGACGGCGTC  239
Gvel_83898_c0_g1      ------------------------------------------------------------  0
Gvel_78482_c1_g1      ------------------------------------------------------------  0
Gvel_78482_c2_g1      ------------------------------------------------------------  0
                      .. ..  . .. ..    ..  . ... .  . ..  . ..  . ..  . .. ..  . 

Gvel_34771_c0_g1      GATTTCTACACTTCTATCACTAGGGCTCGCTTTGAAGAATTGAATGCTGACCTCTTCCGC  1197
Gvel_19946_c0_g1      ------------------------------------------------------------  734
Gvel_16103_c0_g1      GACTTCTACACCAAGGTGACGCGCGCGCGCTTCGAGCAGCTCTGCGACGACCTCTTCCGC  299
Gvel_83898_c0_g1      ------------------------------------------------------------  0
Gvel_78482_c1_g1      ------------------------------------------------------------  0
Gvel_78482_c2_g1      ------------------------------------------------------------  0
                      .. ........     . ..  . .. ..... ..  .  .         ..........
```

**Figure S3 (5/9)**

```
Gvel_34771_c0_g1        AGCACCATGGAACCTGTTGAGAAGTCTTTGCGTGACGCAAAGATCGACAAGGCTCAAATT  1257
Gvel_19946_c0_g1        ------------------------------------------------------------  734
Gvel_16103_c0_g1        TCCACGCTCGACCCCGTGCGCAAGGCGCTGCTCGACGCCAAGGTGGCCACGGCCGACATC  359
Gvel_83898_c0_g1        ------------------------------------------------------------  0
Gvel_78482_c1_g1        ------------------------------------------------------------  0
Gvel_78482_c2_g1        ------------------------------------------------------------  0
                          ...  . .. .. ..    ... .  ...  ..... ... . . .. ...  . ..

Gvel_34771_c0_g1        CACGACATTGTTCTTGTTGGTGGTTCTACTCGCATTCCTAAGGTTCAAAAGCTCCTACAA  1317
Gvel_19946_c0_g1        ------------------------------------------------------------  734
Gvel_16103_c0_g1        GACGAGGTGCTGCTGGTGGGCGGCAGCACGCGCATCCCCAAGATCCAGCGCCTGCTGCAA  419
Gvel_83898_c0_g1        ------------------------------------------------------------  0
Gvel_78482_c1_g1        ----------------------------CGCGCATCCCCAAGATACAGAAGATGCTGCAG  32
Gvel_78482_c2_g1        ------------------------------------------------------------  0
                         ....  .  . .. .. .. ..    .:.::::::.::.::::.: ::......:.::.::.

Gvel_34771_c0_g1        GATTTCTTCAATGGCAAAGAATTGAACAAATCCATCAACCCTGATGAAGCTGTTGCTTAT  1377
Gvel_19946_c0_g1        ------------------------------------------------------------  734
Gvel_16103_c0_g1        GACTTCTTCCACGGCAAGGCGCTCAACCTGTCCATCAACCCGGACGAGGCGGTGGCGTAC  479
Gvel_83898_c0_g1        ------------------------------------------------------------  0
Gvel_78482_c1_g1        GACTTCTTCTGCGGGAAGGCGCTCAACCTGTCCATCAACCCGGACGAGGCGGTGGCGTAC  92
Gvel_78482_c2_g1        ------------------------------------------------------------  0
                        ::.:::::: ..::.::.:...:.:::...:::::::::::.::.::.::.::.::.

Gvel_34771_c0_g1        GGTGCTGCTGTGCAGGCTGCAATTTTGGCTGGTGATAAGTCTGAGGAGGTTCAAGATCTC  1437
Gvel_19946_c0_g1        ------------------------------------------------------------  734
Gvel_16103_c0_g1        GGCGCCGCCGTGCAGGCGGCCGTGC-----------------------------------  504
Gvel_83898_c0_g1        ------------------------------------------------------------  0
Gvel_78482_c1_g1        GGCGCCGCCGTGCAGGCCGCCATCCTGAGCGGCGACACCAGCTCCCAGATCCAGGACGTG  152
Gvel_78482_c2_g1        ------------------------------------------------------------  0
                        ::.::.::.::::::::: ::..: ...    .. .. .          .. . .. ..   .

Gvel_34771_c0_g1        CTTTTGCTTGATGTTACACCATTGTCTCTGGGTATTGAAACTGCTGGAGGAGTCATGACA  1497
Gvel_19946_c0_g1        ------------------------------------------------------------  734
Gvel_16103_c0_g1        ------------------------------------------------------------  504
Gvel_83898_c0_g1        --------------------------------------------------------GACG  4
Gvel_78482_c1_g1        CTGCTGGTGGACGTGGCGCCGCTGTCGCTGGGCATCGAGACGGCCGGCGGCGTCATGAAC  212
Gvel_78482_c2_g1        ------------------------------------------------------------  0
                        ..   .. . .. ..   . ..  .... ..... .. .. .. .. .. .. .....::.
```

**Figure S3 (6/9)**

```
Gvel_34771_c0_g1      GCCCTCATCAAGCGAAACACAACCATTCCAACCAAGCAGACCCAAACTTTCACCACATAT   1557
Gvel_19946_c0_g1      ------------------------------------------------------------   734
Gvel_16103_c0_g1      ------------------------------------------------------------   504
Gvel_83898_c0_g1      AAGATCGTGGAGCGCAACTCGCGCATCCCCTGCAAGCAGACGCAGACCTTCACCACCTAC   64
Gvel_78482_c1_g1      AACATCGTCCACCGCAACACGCGCATCCCGTGCAAGCAGTCGCAGACCTTCACCACCTAC   272
Gvel_78482_c2_g1      -----------------------------------------------CTTCACCACCTAC   13
                      ....::.:  :.::.:::.:...:::.:: ..:::::::.:.::.:::********:**:

Gvel_34771_c0_g1      TCTGATAACCAACCTGGTGTACTCATCCAGGTTTACGAGGGTGAAAGAGCCATGACCAAG   1617
Gvel_19946_c0_g1      ------------------------------------------------------------   734
Gvel_16103_c0_g1      ------------------------------------------------------------   504
Gvel_83898_c0_g1      TCCGACAACCAGCCGGCCGTCACCATCCAGGTGTTCGAGGGCGAGCGCGCCATGACGCGC   124
Gvel_78482_c1_g1      CACGACAACCAAACCGC-------------------------------------------   289
Gvel_78482_c2_g1      CACGACAACCAGACCGCCGTCACCATCCAGGTGTTCGAAGGGGAGCGCGCCATGACGCGC   73
                      ..:**:*****..*.*:.::...:::::::::.:.:::.:: ::..:.::::::::....

Gvel_34771_c0_g1      GACAACAACCTTCTTGGAAAGTTTGAACTGACTGGCATTCCCCCAGCTCCACGTGGTGTC   1677
Gvel_19946_c0_g1      ------------------------------------------------------------   734
Gvel_16103_c0_g1      ------------------------------------------------------------   504
Gvel_83898_c0_g1      GACAACAACCTGCTGGGCACCTTCAACCTGACGGGCATCCCGCCCGCGCCGCGCGGCGTG   184
Gvel_78482_c1_g1      ------------------------------------------------------------   289
Gvel_78482_c2_g1      AACAACAACCTGCTGGGCACGTTCGACCTGACGGGGCTGGCGGCGGCGCCGCGCGGCGCG   133
                      .::::::::::.::.::.:. ::..:.:::::.::..: .:..: ::.::.::.::.: .

Gvel_34771_c0_g1      CCACAAATTGAAGTTACTTTTGATATTGATGCTAATGGTATTTTGAATGTAACTGCTATT   1737
Gvel_19946_c0_g1      ------------------------------------------------------------   734
Gvel_16103_c0_g1      ------------------------------------------------------------   504
Gvel_83898_c0_g1      CCCAAGATCGAGGTGACGTTCGACCTGGACGCGAACGGCATCCTCAACGTGTCCGCCAAG   244
Gvel_78482_c1_g1      ------------------------------------------------------------   289
Gvel_78482_c2_g1      CCCAAGATCGACGTCACGTTCGACCTGGACGCCGACGGCATCCTCAACGTGTCGGCGACG   193
                      ::..:.::.:: :: ::.::.::..:.::.:: .:.::.::..:.::.::..: :: : .

Gvel_34771_c0_g1      GAGAAATCTACTGGAAAAGAAAACAAGATCACAATCACCAATGACAAGGGTCGTCTTAGC   1797
Gvel_19946_c0_g1      ------------------------------------------------------------   734
Gvel_16103_c0_g1      ------------------------------------------------------------   504
Gvel_83898_c0_g1      GACAGCAGCACGGGCAAGTCGGAGCGCATCACCATCCAGAACGACAAGGGCCGCCTCTCC   304
Gvel_78482_c1_g1      ------------------------------------------------------------   289
Gvel_78482_c2_g1      GACGCCAGCTCGGGGCGGCGGCGCGCCGTCACCATCCGCAACGAGCGCGGCCGCCTCTCC   253
                      ::.. .....:.:: ...  . ..  ..::::.:::.  ::.::....::.::.::...:
```

**Figure S3 (7/9)**

```
Gvel_34771_c0_g1     AAGGAAGAAATAGAGCGTATGGTTAATGATGCTGAAAAATACCGTGCTGAAGATGACAAA  1857
Gvel_19946_c0_g1     ------------------------------------------------------------  734
Gvel_16103_c0_g1     ------------------------------------------------------------  504
Gvel_83898_c0_g1     AAGGACGAGATCGACCGCATGCTGTCGGAGGCCGAGCGCTACCGCGCCGAGGACGAGAAG  364
Gvel_78482_c1_g1     ------------------------------------------------------------  289
Gvel_78482_c2_g1     AGCGACGACATCGCGCGCATGCTGGCCGACGCGGAGCGCTACCGCGGCGAGAA-------  306
                     :...:.:: ::.: .::.:::.:. . :: :: ::....:::::.:..::..: .. ..

Gvel_34771_c0_g1     CAGAAACAGACCATTGCTGCTAAGAATTCCTTGGAATCCTACTGTTTCAACATGAAGAGC  1917
Gvel_19946_c0_g1     ------------------------------------------------------------  734
Gvel_16103_c0_g1     ------------------------------------------------------------  504
Gvel_83898_c0_g1     CAGCGCGAGCGCGTCGCCGCGCGCAACCAGCTCGAGAGCTACACCTTCAGCGTCAAGCAG  424
Gvel_78482_c1_g1     ------------------------------------------------------------  289
Gvel_78482_c2_g1     ------------------------------------------------------------  306
                     ...    ..  . . .. ..    ..     . ..   ....   .... . . ...

Gvel_34771_c0_g1     ACTGTTGAAGACGAGAAGCTGAAGGATAAGATCTCTGATGCTGATAAGACTTCAATTTTG  1977
Gvel_19946_c0_g1     ------------------------------------------------------------  734
Gvel_16103_c0_g1     ------------------------------------------------------------  504
Gvel_83898_c0_g1     GCCACCGAAGACGCGC---CGGCGGGCAAGCTGACGGACGCCGACAAGAAGACGGTGCAG  481
Gvel_78482_c1_g1     ------------------------------------------------------------  289
Gvel_78482_c2_g1     ------------------------------------------------------------  306
                      .    ....... .     .  ..  ... .  . .. .. .. ....   .  .   .

Gvel_34771_c0_g1     GACAAGTGCAATGAGGTCATTCGCTGGCTGGATTCTAACCAACTTGCTGAAAAAGAAGAA  2037
Gvel_19946_c0_g1     ------------------------------------------------------------  734
Gvel_16103_c0_g1     ------------------------------------------------------------  504
Gvel_83898_c0_g1     GAGCGCTGCGCCGCCGTCCTCTCGTGGCTGGACGGCAACACCTTGGCCGAGAAGGACGAG  541
Gvel_78482_c1_g1     ------------------------------------------------------------  289
Gvel_78482_c2_g1     ------------------------------------------------------------  306
                     ..    ...   .  ... .    ........    ...    . .. .. .. .. ..

Gvel_34771_c0_g1     TTCGAGGCCCAACAGAAGGAACTTGAAGCGCTCTGCAACCCAATAATCACCAAGTTGTAC  2097
Gvel_19946_c0_g1     ------------------------------------------------------------  734
Gvel_16103_c0_g1     ------------------------------------------------------------  504
Gvel_83898_c0_g1     TACGAGGCGAAGCTGAAGGAGCTGCAGCGCGACTGCGCGCCCGTCATGACCAAGCTGCAC  601
Gvel_78482_c1_g1     ------------------------------------------------------------  289
Gvel_78482_c2_g1     ------------------------------------------------------------  306
                     . ......  . . ...... ..  .      ....   ..  . .. ...... .. ..
```

**Figure S3 (8/9)**

```
Gvel_34771_c0_g1     CAGGGTGCTGGTGGAGCTCCTGGTGGCATGCCTGGATTTCCTGGTGGTTTCCCTGGTGGC   2157
Gvel_19946_c0_g1     ------------------------------------------------------------   734
Gvel_16103_c0_g1     ------------------------------------------------------------   504
Gvel_83898_c0_g1     CAGGGCGCGGCGGGCCAGGGCGGCGCGCCGGGC----GCCAAGGGCCCCACCGTGGAGGA   657
Gvel_78482_c1_g1     ------------------------------------------------------------   289
Gvel_78482_c2_g1     ------------------------------------------------------------   306
                     ..... .. .  ..       .. .    .            ..         ... ..

Gvel_34771_c0_g1     GGTGCCGCTCCTGGACCTG-GTGGTGCTGCTGGTGCTGGAGCCGGTGCTGGACCTACTAT   2216
Gvel_19946_c0_g1     ------------------------------------------------------------   734
Gvel_16103_c0_g1     ------------------------------------------------------------   504
Gvel_83898_c0_g1     GGTGGACTAGACGCCCCGCCCTGCCCCGCCCCGCGCCGCCCCCGCCCCCGCCCCCGACAC   717
Gvel_78482_c1_g1     ------------------------------------------------------------   289
Gvel_78482_c2_g1     ------------------------------------------------------------   306
                     ....           ..    ..      .  . .. .   ...   . .  ..    .

Gvel_34771_c0_g1     TGAAGAAGTTGATTAAACAATTCATCACTTGCAACCATTTTAGTTCATGGCTAAAGTTTC   2276
Gvel_19946_c0_g1     ------------------------------------------------------------   734
Gvel_16103_c0_g1     ------------------------------------------------------------   504
Gvel_83898_c0_g1     GCACGCGGGACGCGCCGGCGCTCGCGAAC-------------------------------   746
Gvel_78482_c1_g1     ------------------------------------------------------------   289
Gvel_78482_c2_g1     ------------------------------------------------------------   306
                       . .  .             ..   .

Gvel_34771_c0_g1     ATTCCAGAATCAGTATTATTGCAGACACAACTTAAAGTCAATTTTCAAAGTGAGATGTCT   2336
Gvel_19946_c0_g1     ------------------------------------------------------------   734
Gvel_16103_c0_g1     ------------------------------------------------------------   504
Gvel_83898_c0_g1     ------------------------------------------------------------   746
Gvel_78482_c1_g1     ------------------------------------------------------------   289
Gvel_78482_c2_g1     ------------------------------------------------------------   306

Gvel_34771_c0_g1     GCTTAAAGCAAGAAACAAGAAAATTTATTGTATTTATTCTGCATCTAAATTCACTTGATT   2396
Gvel_19946_c0_g1     ------------------------------------------------------------   734
Gvel_16103_c0_g1     ------------------------------------------------------------   504
Gvel_83898_c0_g1     ------------------------------------------------------------   746
Gvel_78482_c1_g1     ------------------------------------------------------------   289
Gvel_78482_c2_g1     ------------------------------------------------------------   306
```

**Figure S3 (9/9)**

```
Gvel_34771_c0_g1        TTTGGCAAACTTGAATGTTCTGTTAAATTTTCTTCAGTTTTTAAAAATTAAAGTAATTGG  2456
Gvel_19946_c0_g1        ------------------------------------------------------------  734
Gvel_16103_c0_g1        ------------------------------------------------------------  504
Gvel_83898_c0_g1        ------------------------------------------------------------  746
Gvel_78482_c1_g1        ------------------------------------------------------------  289
Gvel_78482_c2_g1        ------------------------------------------------------------  306

Gvel_34771_c0_g1        CATACAGAAAAAAA  2470
Gvel_19946_c0_g1        --------------  734
Gvel_16103_c0_g1        --------------  504
Gvel_83898_c0_g1        --------------  746
Gvel_78482_c1_g1        --------------  289
Gvel_78482_c2_g1        --------------  306
```

**Figure S3 (above)**. Alignment of putative *Gryllus veletis* HSP70 transcripts from Toxopeus et al. 2019a that had BLAST hits to other Orthoptera (Table S1). Arrows above sequence indicate the forward and reverse primers used in RT-qPCR. Produced using Clustal Omega, and modified with the following symbols: one dot (.) indicates the same nucleotide in two sequences, two dots (:) indicates the same nucleotide in two sequences, and an asterisk (*) indicates the same nucleotide in four sequences.

**Figure S4 (1/8)**

```
S.americana.Hsp70A1    CGAAGTTAGTGCTTCGAGACGTGATCGAAGATTCGCTACGCAGCTATATTCGTCTACGTT    180
S.americana.HSP70B2    ------------------------------------GGCACAGCAGCAGCCAGCAGCCAG    24
S.serialis.HSP70B2     ----------------------------------------CAGCAGCAGCCAGCAGCCAG    20
Gvel_19946_c0_g1       -----------------------------------GCGGAGGACGCGGCGCCCGAGAGCG    25
Gvel_16103_c0_g1       ------------------------------------------------------------    0
Gvel_83898_c0_g1       ------------------------------------------------------------    0
                                                            ...:::*...:..*..::::..:

S.americana.Hsp70A1    ACTGTGACATAGCTAACAGAAGAAAATGCCGAAGATTCCAGCAGTGGGGATCGATTTGGG    240
S.americana.HSP70B2    CAACCAGCAGCCATCAGCAGCGACCATGGGCAAGGCAACCGCCGTCGGCATAGACCTGGG    84
S.serialis.HSP70B2     CAACCAGCAGCCATCAGCAGCGACCATGGGCAAGGCAACCGCCGTCGGCATAGACCTGGG    80
Gvel_19946_c0_g1       ACATTTGAACAGCATTTCAGCCACCATGGGGAAAGTGCCTGCAGTGGGAATTGATCTCGG    85
Gvel_16103_c0_g1       ------------------------------------------------------------    0
Gvel_83898_c0_g1       ------------------------------------------------------------    0
                       ..:...::*....:.:.:::::*::***::.**::...*.**.**.**.**.**.:*:**

S.americana.Hsp70A1    AACCACGTACTCGTGCGTGGGAGTGTGGCAACAAGGCAAGGTGGAAATAATCGCCAACGA    300
S.americana.HSP70B2    CACCACCTACTCGTGCGTGGGCGTGTGGCAGCACGGCAAGGTCGAGATCGTCGCCAACGA    144
S.serialis.HSP70B2     CACCACCTACTCGTGCGTGGGCGTGTGGCAGCACGGCAAGGTCGAGATCGTCGCCAACGA    140
Gvel_19946_c0_g1       CACGACCTATTCGTGCGTCGGGGTCTGGCAGCATGGTAAAGTAGAGATAATCGCCAATGA    145
Gvel_16103_c0_g1       ------------------------------------------------------------    0
Gvel_83898_c0_g1       ------------------------------------------------------------    0
                       :**:**:**:********:**.**:*****:**.**:**:*.**:***..*******:**

S.americana.Hsp70A1    TCAAGGCAACAGGACGACACCAAGCTACGTCGCCTTTTGCGACTCGGAGCGGCTGATCGG    360
S.americana.HSP70B2    GCAGGGCAACCGCACCACGCCCAGCTACGTCGCCTTCACCGACACCGAGCGACTCATCGG    204
S.serialis.HSP70B2     GCAGGGCAACCGCACCACGCCCAGCTACGTCGCCTTCACCGACACCGAACGACTCATCGG    200
Gvel_19946_c0_g1       CCAAGGAAATCGCACAACACCCAGTTATGTCGCTTTCACGGACACTGAACGGCTTATCGG    205
Gvel_16103_c0_g1       ------------------------------------------------------------    0
Gvel_83898_c0_g1       ------------------------------------------------------------    0
                       .**.**:**::*:**.**.**:**:**:*****:**::::***:*.**.**.**.*****

S.americana.Hsp70A1    CGATGCTGCCAAGAACCAGGTTGCGATGAACCCGCAGAATACCATCTTCGACGCCAAACG    420
S.americana.HSP70B2    CGATGCCGCCAAAAGCCAGGTGGCCATGAACCCGAAGAACACGGTTTTCGACGCGAAGCG    264
S.serialis.HSP70B2     CGATGCCGCCAAGAGCCAGGTGGCCATGAACCCGAAGAACACGGTTTTCGACGCGAAGCG    260
Gvel_19946_c0_g1       CGATGCTGCCAAAAGTCAGGTGGCGATGAACCCAAAGAACACCGTTTTCGACGCCAAACG    265
Gvel_16103_c0_g1       ------------------------------------------------------------    0
Gvel_83898_c0_g1       ------------------------------------------------------------    0
                       ******.*****.*::*****:**.********::****:**.:*:********.**.**
```

**Figure S4 (2/8)**

```
S.americana.Hsp70A1     ACTCATCGGCCGCAAGTTTGACGACCCAAAGGTGCAGGGAGACATGAAGCATTGGCCATT    480
S.americana.HSP70B2     ATTGATTGGAAGACGCTTTGACGATCCCAAGATACAAGACGACATGAAGCACTGGCCGTT    324
S.serialis.HSP70B2      ATTGATTGGAAGACGCTTTGACGATCCCAAGATACAAGACGACATGAAGCACTGGCCGTT    320
Gvel_19946_c0_g1        CTTAATAGGAAGGCGCTTCGATGATGCGAAGATTCAAGACGACATGAAGCACTGGCCGTT    325
Gvel_16103_c0_g1        ------------------------------------------------------------    0
Gvel_83898_c0_g1        ------------------------------------------------------------    0
                        ::*.**.**::*.:::**:**:**::*.***:*.**:*::***********:*****:**

S.americana.Hsp70A1     CAAAGTGATTAACGACTGTAGCAAGCCTAAGATACAGGTCCAGTTCAAAGGCACCACCAA    540
S.americana.HSP70B2     CACGGTCGTTTCCGACGGCGACAAACCCAAAATTCAGGTGGAGTACAAGGGCGAGACGAA    384
S.serialis.HSP70B2      CACGGTCGTTTCCGACGGCGACAAACCCAAAATTCAGGTGGAGTACAAGGGCGAGACGAA    380
Gvel_19946_c0_g1        TACTGTCGTCAACGATGGCGGCAAGCCCAAGATCCAGGTGGAGTACAAGAGCGAGGTGAA    385
Gvel_16103_c0_g1        ------------------------------------------------------------    0
Gvel_83898_c0_g1        ------------------------------------------------------------    0
                        :*:.**::*:..***::*::.***.**:**.**.*****::***:***::**:::::**

S.americana.Hsp70A1     AACTTTCGCTCCAGAAGAGATAAGCGCTATGGTGCTTGTGAAAATGAAGGAAACAGCAGA    600
S.americana.HSP70B2     GAGGTTCGCGCCCGAGGAGATCAGCTCGATGGTGCTGAGCAAGATGCGCGAGATCGCGAG    444
S.serialis.HSP70B2      GAGGTTCGCGCCCGAGGAGATCAGCTCGATGGTGCTGAGCAAGATGCGCGAGATCGCGAG    440
Gvel_19946_c0_g1        GCGCTTCGCGCCGGAGGAGGTGAGCTCCATGGTGCTGACCAAGATGCGCGAGATCGCCGA    445
Gvel_16103_c0_g1        ------------------------------------------------------------    0
Gvel_83898_c0_g1        ------------------------------------------------------------    0
                        :::.*****:**.**:***:*.***:*.********::.:**:***:::**:*::**...

S.americana.Hsp70A1     GGCATTCTTGGGTGGGCAGGTTCGAGAGGCAGTTATTACAGTGCCCGCCTACTTCAACGA    660
S.americana.HSP70B2     CACGTACCTGGGCGGCGACGTGCGCGACGCGGTGATCACGGTGCCGGCCTACTTCAACGA    504
S.serialis.HSP70B2      CACGTACCTGGGCGGCGACGTGCGCGACGCGGTGATCACGGTGCCGGCCTACTTCAACGA    500
Gvel_19946_c0_g1        GACGTACCTGGGCGGGAAGGTGACGGACGCGGTGATCACGGTGCCGGCCTACTTCAACGA    505
Gvel_16103_c0_g1        ------------------------------------------------------------    0
Gvel_83898_c0_g1        ------------------------------------------------------------    0
                        .:*:*:*:****:**..*.**:::.**:**:**:**:**:*****:*************

S.americana.Hsp70A1     CTCACAGCGGCAAGCTACCAAAGATGCAGGCGCCATAGCAGGCCTGAAGGTGCTGCGGAT    720
S.americana.HSP70B2     CGCGCAGCGCAGCGCCACCAAGGACGCGGGCGCCATCGCCGGCCTCAACGTGCTGCGCAT    564
S.serialis.HSP70B2      CGCGCAGCGCAGCGCCACCAAGGACGCGGGCGCCATCGCCGGCCTCAACGTGCTGCGCAT    560
Gvel_19946_c0_g1        CTCGCAGCGGCAGGCCACCAAGGACGCCGGCGCCATCGCGGGCCTCAACGTGCTGCGGAT    565
Gvel_16103_c0_g1        ------------------------------------------------------------    0
Gvel_83898_c0_g1        ------------------------------------------------------------    0
                        *.*:*****....**:*****:**:**.********:**.*****:**:********.**
```

**Figure S4 (3/8)**

```
S.americana.Hsp70A1    TATCAACGAGCCAACTGCAGCTGCACTCGCATACGGCCTCGACAAAAACCTGAAAGGTGA    780
S.americana.HSP70B2    CATCAACGAGCCCACCGCCGCCGCGCTCGCCTACGGGCTCGACAAGAACCTGCAGGGCGA    624
S.serialis.HSP70B2     CATCAACGAGCCCACCGCCGCCGCGCTCGCCTACGGGCTCGACAAGAACCTGCAGGGCGA    620
Gvel_19946_c0_g1       CATCAACGAGCCGACGGCCGCCGCGCTGGCCTACGGCCTCGACAAGAACCTCAAGGGCGA    625
Gvel_16103_c0_g1       ------------------------------------------------------------    0
Gvel_83898_c0_g1       ------------------------------------------------------------    0
                       :***********.**.**:**:**:**:**:*****.********:*****:.*:**:**

S.americana.Hsp70A1    GCGCAATGTGCTCATATTTGACCTTGGTGGTGGCACCTTTGACGTGTCCATCCTGAGCAT    840
S.americana.HSP70B2    GAAGAACGTGCTCATCTTCGACCTCGGCGGAGGCACTTTTGACGTGTCGGTGCTGGCCAT    684
S.serialis.HSP70B2     GAAGAACGTGCTCATCTTCGACCTCGGCGGAGGCACTTTTGACGTGTCGGTGCTGGCCAT    680
Gvel_19946_c0_g1       GAAGAACGTGCTCATCTTCGACCTCGGCGGCGGCACCTTCGACGTCTCCATCCTCTCCAT    685
Gvel_16103_c0_g1       ------------------------------------------------------------    0
Gvel_83898_c0_g1       ------------------------------------------------------------    0
                       *:::**:********:**:*****:**:**.*****.**:*****:**..*.**:.:***

S.americana.Hsp70A1    TTCGGAGGGTTCACTGTTTGAAGTGAAAGCAACAGCTGGAGACACACACCTGGGAGGCGA    900
S.americana.HSP70B2    CTCGGAAGGGTCGCTGTTCGAGGTGAAGTCGACGGCGGGTGATACGCACCTGGGCGGCGA    744
S.serialis.HSP70B2     CTCGGAAGGGTCGCTGTTCGAGGTGAAGTCGACGGCGGGTGATACGCACCTGGGCGGCGA    740
Gvel_19946_c0_g1       CGACGAGGGATCGCTCTTCGAGGTGAAGTCGACGGCCGGCGACACGCAC-----------    734
Gvel_16103_c0_g1       --CCGAGGGCTCGCTCTTCGAGGTGCGGGCGACGGCCGGCGACACGCACCTGGGCGGCGA    58
Gvel_83898_c0_g1       ------------------------------------------------------------    0
                       ::*:**:**.*****:************:*******.**.**:***********:*****

S.americana.Hsp70A1    GGACTTTGACAGTCGGCTTGTGAATCATTTGGCTGATGAGTTCAAACGTAAATTCCACAA    960
S.americana.HSP70B2    GGACTTCGACAACCGGCTGGTGCAGCACCTGGCGGAAGAGTTCCAGCGCAAGCACCGCAA    804
S.serialis.HSP70B2     GGACTTCGACAACCGGCTGGTGCAGCACCTGGCAGAGGAGTTCCAGCGCAAGCACCGCAA    800
Gvel_19946_c0_g1       ------------------------------------------------------------    734
Gvel_16103_c0_g1       GGACTTCGACACGCGCCTCGTCGCCCACCTCGCCGACGAGTTCCGGCGCAAGCACGGCAA    118
Gvel_83898_c0_g1       ------------------------------------------------------------    0
                       ******:****..**:**.**:.:.**::*:** ** ******:::**:**:::*::***

S.americana.Hsp70A1    GGACGTACGTTCCAATCCACGTGCTTTGCGTCGACTGCGTACGGCAGCAGAACGGGCCAA    1020
S.americana.HSP70B2    GGACATGCGCGCCAACGCGCGCGCGCTGCGCCGCCTGCGCACCGCCGCCGAGCGCGCCAA    864
S.serialis.HSP70B2     GGACATGCGCGCCAACGCGCGCGCGCTGCGCCGCCTGCGCACCGCCGCCGAGCGCGCCAA    860
Gvel_19946_c0_g1       ------------------------------------------------------------    734
Gvel_16103_c0_g1       GGACGTGCGCCGCCACGCGCGCGCCCTGCGCCGCCTGCGCACCGCCGCCGAGCGCGCCAA    178
Gvel_83898_c0_g1       ------------------------------------------------------------    0
                       ****.*:**:.:*:*::*:**:**.:****:**:*****:**:**:**:**:**:*****
```

**Figure S4 (4/8)**

```
S.americana.Hsp70A1     GCGTACGTTGTCTTCCAGCACGGAGGCCAGTATTGAGATTGATGCACTATTAGATGGCAT    1080
S.americana.HSP70B2     GCGGACGCTCTCGTCCAGCACCGAGGCCAGCCTCGAGATAGACGCCCTGCACGACGGCAT    924
S.serialis.HSP70B2      GCGGACGCTCTCGTCCAGCACCGAGGCCAGCCTCGAGATAGACGCCCTGCACGACGGCAT    920
Gvel_19946_c0_g1        ------------------------------------------------------------    734
Gvel_16103_c0_g1        GCGCACCCTCTCCTCCAGCACCGAGGCCGGCGTCGAGGTGGACGCCCTGCTCGACGGCGT    238
Gvel_83898_c0_g1        ------------------------------------------------------------    0
                        ***:**::*:**.********:******:*:.*:***:*.**:**:**::.:**:***:*

S.americana.Hsp70A1     TGACTTCTACACAAAAGTTTCACGAGCACGGTTTGAGGAACTTTGTGCTGATCTGTTCCG    1140
S.americana.HSP70B2     CGACTTCTACGCCAAGGTGACGCGCGCGCGGTTCGAGGAGCTGTGCATGGACCTCTTCCG    984
S.serialis.HSP70B2      CGACTTCTACGCCAAGGTGACGCGCGCGCGGTTCGAGGAGCTGTGCATGGACCTCTTCCG    980
Gvel_19946_c0_g1        ------------------------------------------------------------    734
Gvel_16103_c0_g1        CGACTTCTACACCAAGGTGACGCGCGCGCGCTTCGAGCAGCTCTGCGACGACCTCTTCCG    298
Gvel_83898_c0_g1        ------------------------------------------------------------    0
                        :*********.*:**:**::*:**:**:**:**:***:*:**.**:...**:**:*****

S.americana.Hsp70A1     CTCAACACTACAACCTGTCGAGAAAGCACTGGCCGATGCCAAGATGGACAAAGCGTCAAT    1200
S.americana.HSP70B2     GCAGACGCTGGCGCCGGTGGAGCGCGCCCTGGGCGACGCCAAGCTGGACAAGGCGTCCGT    1044
S.serialis.HSP70B2      GCAGACGCTGGCGCCGGTGGAGCGCGCCCTGGGCGACGCCAAGCTGGACAAGGCGTCCGT    1040
Gvel_19946_c0_g1        ------------------------------------------------------------    734
Gvel_16103_c0_g1        CTCCACGCTCGACCCCGTGCGCAAGGCGCTGCTCGACGCCAAGGTGGCCACGGCCGACAT    358
Gvel_83898_c0_g1        ------------------------------------------------------------    0
                        ....**:**.:..**.**:::...**.***:.***:******.***:**::**:*::.*

S.americana.Hsp70A1     CCACGATGTCGTGTTGGTGGGTGGTTCAACACGCATCCCTAAGATTCAGGCTCTGCTGCA    1260
S.americana.HSP70B2     GCACGACGTCGTGCTGGTCGGCGGCTCCACGCGCATCCCCAAGATACAGAAGATGCTGCA    1104
S.serialis.HSP70B2      GCACGACGTCGTGCTGGTCGGCGGCTCCACGCGCATCCCCAAGATACAGAAGATGCTGCA    1100
Gvel_19946_c0_g1        ------------------------------------------------------------    734
Gvel_16103_c0_g1        CGACGAGGTGCTGCTGGTGGGCGGCAGCACGCGCATCCCCAAGATCCAGCGCCTGCTGCA    418
Gvel_83898_c0_g1        ------------------------------------------------------------    0
                        .:****.**::**:****:**:**::::**:*******:*****.***....*******

S.americana.Hsp70A1     GAACTACTTTGCTGGTAAGCGGCTGAATTTGTCCATCAACCCGGATGAGGCAGTGGCATA    1320
S.americana.HSP70B2     GGACTTCTTCTGCGGCAAGACGCTGAACCTGTCCATCAACCCGGACGAGGCGGTGGCGTA    1164
S.serialis.HSP70B2      GGACTTCTTCTGCGGCAAGACGCTGAACCTGTCCATCAACCCGGACGAGGCGGTGGCGTA    1160
Gvel_19946_c0_g1        ------------------------------------------------------------    734
Gvel_16103_c0_g1        AGACTTCTTCCACGGCAAGGCGCTCAACCTGTCCATCAACCCGGACGAGGCGGTGGCGTA    478
Gvel_83898_c0_g1        ------------------------------------------------------------    0
                        ::***:***:..:**:***.:***:**::***************:*****:*****:**
```

**Figure S4 (5/8)**

```
S.americana.Hsp70A1    TGGTGCAGCTGTACAAGCAGCCATTCTCAGTGGTGATACCAGCTCTGCAATTCAGGATGT    1380
S.americana.HSP70B2    CGGCGCGGCGGTGCAGGCGGCCATCCTGAGCGGCGACACGAGCTCGCAGATCCAGGACGT    1224
S.serialis.HSP70B2     CGGCGCGGCGGTGCAGGCGGCCATCCTGAGCGGCGACACGAGCTCGCAGATCCAGGACGT    1220
Gvel_19946_c0_g1       ------------------------------------------------------------    734
Gvel_16103_c0_g1       CGGCGCCGCCGTGCAGGCGGCCGTGC----------------------------------    504
Gvel_83898_c0_g1       ------------------------------------------------------------    0
                       :**:**.**.**:**:**:***:*.*:.::.::.::.::.:::::....::.:::::.::

S.americana.Hsp70A1    GTTACTGGTAGATGTAGCACCACTATCACTCGGCATTGAGACAGCAGGTGGTGTCATGAC    1440
S.americana.HSP70B2    GCTGCTGGTCGACGTGGCGCCGCTCTCGCTCGGCATCGAGACCGCGGGCGGCGTCATGAC    1284
S.serialis.HSP70B2     GCTGCTGGTCGACGTGGCGCCGCTCTCGCTCGGCATCGAGACCGCGGGCGGCGTCATGAC    1280
Gvel_19946_c0_g1       ------------------------------------------------------------    734
Gvel_16103_c0_g1       ------------------------------------------------------------    504
Gvel_83898_c0_g1       ---------------------------------------------------------GAC    3
                       :.:.:::::.::.::.::.::.::.::.::::::::.:::::.::.::.::.:::::***

S.americana.Hsp70A1    TAAGCTGGTAGAGCGCAATGCGCGTATCCCCTGCAAGCAAAAACAGACATTCACCACATA    1500
S.americana.HSP70B2    CAAGATCGTCGAGCGCAACGCGCGCATCCCCTGCAAGCAGAAGCAGACCTTCACCACCTA    1344
S.serialis.HSP70B2     CAAGATCGTCGAGCGCAACGCGCGCATCCCCTGCAAGCAGAAGCAGACCTTCACCACCTA    1340
Gvel_19946_c0_g1       ------------------------------------------------------------    734
Gvel_16103_c0_g1       ------------------------------------------------------------    504
Gvel_83898_c0_g1       GAAGATCGTGGAGCGCAACTCGCGCATCCCCTGCAAGCAGACGCAGACCTTCACCACCTA    63
                       .***:*:**.********::****:**************:*::*****:********:**

S.americana.Hsp70A1    CTCAGACAACCAGCCAGCAGTTACAATTCAGGTGTTTGAAGGAGAGCGTGCAATGACAAA    1560
S.americana.HSP70B2    CTCCGACAACCAGCCCGCCGTCACCATTCAGGTGTTCGAGGGCGAGCGCGCCATGACCAA    1404
S.serialis.HSP70B2     CTCCGACAACCAGCCCGCCGTCACCATTCAGGTGTTCGAGGGCGAGCGCGCCATGACCAA    1400
Gvel_19946_c0_g1       ------------------------------------------------------------    734
Gvel_16103_c0_g1       ------------------------------------------------------------    504
Gvel_83898_c0_g1       CTCCGACAACCAGCCGGCCGTCACCATCCAGGTGTTCGAGGGCGAGCGCGCCATGACGCG    123
                       ***:***********.**:**:**:***********:**:**:*****:**:*****.::

S.americana.Hsp70A1    GGACAACAACTTGCTGGGCACATTTGACCTCACTGGCATTCCACCAGCACCACGTGGTGT    1620
S.americana.HSP70B2    GGACAACAACCTGCTGGGCACGTTCAACCTGACGGGCATCCCGCCGGCGCCGCGGGGAGT    1464
S.serialis.HSP70B2     GGACAACAACCTGCTGGGCACGTTCAACCTGACGGGCATCCCGCCGGCGCCGCGGGGAGT    1460
Gvel_19946_c0_g1       ------------------------------------------------------------    734
Gvel_16103_c0_g1       ------------------------------------------------------------    504
Gvel_83898_c0_g1       CGACAACAACCTGCTGGGCACCTTCAACCTGACGGGCATCCCGCCCGCGCCGCGCGGCGT    183
                       :*********:**********.**::****:**:*****:**:**:**:**:**.**.**
```

**Figure S4 (6/8)**

```
S.americana.Hsp70A1     GCCTCAAGTGGAAGTTACATTTGACTTGAATGCAGATGGCATTCTAACTGTGTCCGCCCA   1680
S.americana.HSP70B2     ACCCAAGATCGAGGTGACCTTCGACCTGGACGCGAACGGCATCCTCAACGTGTCGGCGAC   1524
S.serialis.HSP70B2      ACCCAAGATCGAGGTGACCTTCGACCTGGACGCGAACGGCATCCTCAACGTGTCGGCGAC   1520
Gvel_19946_c0_g1        ------------------------------------------------------------   734
Gvel_16103_c0_g1        ------------------------------------------------------------   504
Gvel_83898_c0_g1        GCCCAAGATCGAGGTGACGTTCGACCTGGACGCGAACGGCATCCTCAACGTGTCCGCCAA   243
                        .**:*:::*:**:**:**.**:***:**:*:**::*:*****:**:*::*****.**.::

S.americana.Hsp70A1     AGAGAAGAGCACAGGAAGATCTCGTAACATCACCATCCGCAATGACAAGGGTCGGCTTTC   1740
S.americana.HSP70B2     GGAAAGCGGCTCGGGTCGCAGCGAGCGCATCACCATCCAGAACGACAAGGGCCGCCTCTC   1584
S.serialis.HSP70B2      GGAGAGCGGCTCGGGTCGCAGCGAGCGCATTACCATCCAGAACGACAAGGGCCGCCTTTC   1580
Gvel_19946_c0_g1        ------------------------------------------------------------   734
Gvel_16103_c0_g1        ------------------------------------------------------------   504
Gvel_83898_c0_g1        GGACAGCAGCACGGGCAAGTCGGAGCGCATCACCATCCAGAACGACAAGGGCCGCCTCTC   303
                        :**.*::.**.*:**..:....:::::***:*******::**:********:**:**:**

S.americana.Hsp70A1     CAAGGAGGAAATCGATCGGATGGTAGCTGATGCAGAGCGCTTCAAGGAGGAAGACCAGCG   1800
S.americana.HSP70B2     CAAGGCCGAGATCGAGCGCATGCTGGCCGACGCAGAGCGCTTCCGCGCCGAGGACGAGCG   1644
S.serialis.HSP70B2      CAAGGCCGAGATCGAGCGCATGCTGGCCGACGCAGAGCGCTTCCGCGCCGAGGACGAGCG   1640
Gvel_19946_c0_g1        ------------------------------------------------------------   734
Gvel_16103_c0_g1        ------------------------------------------------------------   504
Gvel_83898_c0_g1        CAAGGACGAGATCGACCGCATGCTGTCGGAGGCCGAGCGCTACCGCGCCGAGGACGAGAA   363
                        *****.:**:*****.**:***:*::*.**.**:*******:*:::*::**:***:**::

S.americana.Hsp70A1     GCAACGTGAGAGAGTAGAGGCACGGCATCGACTGGAGGGCTATGCCCTGTCTGTGAAGCA   1860
S.americana.HSP70B2     GCAGCGCGCGCGGGTCGAGGCCCGCAACCGCCTGGAAGCGTACGCGCTCTCGCTCAAGCA   1704
S.serialis.HSP70B2      GCAGCGCGCGCGGGTCGAGGCCCGCAACCGCCTGGAAGCGTACGCGCTCTCGCTCAAGCA   1700
Gvel_19946_c0_g1        ------------------------------------------------------------   734
Gvel_16103_c0_g1        ------------------------------------------------------------   504
Gvel_83898_c0_g1        GCAGCGCGAGCGCGTCGCCGCGCGCAACCAGCTCGAGAGCTACACCTTCAGCGTCAAGCA   423
                        ***:**:*.*:*.**:*::**.**::*:*:.**:**.:..**::*.:*:::..*:*****

S.americana.Hsp70A1     GGCACTGTCGGACGC---AGGCAGCAGGCTATCTGAGTCGGACAAGACAGCAGCGATGAC   1917
S.americana.HSP70B2     GGCCGCCGAGGACGC---GGGCAGCAAGCTGAGCGACGCGGACAAGGCGACGGTGCGGGA   1761
S.serialis.HSP70B2      GGCCGCCGAGGACGC---GGGCAGCAAGCTGAGCGACGCGGACAAGGCGACGGTGCGGGA   1757
Gvel_19946_c0_g1        ------------------------------------------------------------   734
Gvel_16103_c0_g1        ------------------------------------------------------------   504
Gvel_83898_c0_g1        GGCCACCGAAGACGCGCCGGCGGGCAAGCTGACGGACGCCGACAAGAAGACGGTGCAGGA   483
                        ***:.:::::*****   :**::***:***::.:**::*:******.:::*:*:*:.*::
```

**Figure S4 (7/8)**

```
S.americana.Hsp70A1      ACGTTGTGAGGAGGAATTACAGTGGCTAGACAACAATGCCCAGGCGAGCTGTGGTGAGTG     1977
S.americana.HSP70B2      CCAGAGCGCGGAGGCGCTCAAGTGGCTGGACAGCAACTCGCTGGCGGAGCAGGAGGAGTT     1821
S.serialis.HSP70B2       CCAGAGCGCGGAGGCGCTCAAGTGGCTGGACAGCAACTCGCTGGCGGAGCAGGAGGAGTT     1817
Gvel_19946_c0_g1         ------------------------------------------------------------     734
Gvel_16103_c0_g1         ------------------------------------------------------------     504
Gvel_83898_c0_g1         GCGCTGCGCCGCCGTCCTCTCGTGGCTGGACGGCAACACCTTGGCCGAGAAGGACGAGTA     543
                         .*...*:*::*::*..:*:.:******:***::***:.*.::***::::.::*:.****.

S.americana.Hsp70A1      TGAGCGTCGGCTACAGGAGCTGCAGCGGGTATGCACACCTATAATGACTAAGCTGCACCA     2037
S.americana.HSP70B2      CGAGGACCGCTACAAGCAGCTGTCGGCCGCCTGCTCGCCCATCATGGCCAAGCTGCACCA     1881
S.serialis.HSP70B2       CGAGGACCGCTACAAGCAGCTGTCGGCCGCCTGCTCGCCCATCATGGCCAAGCTGCACCA     1877
Gvel_19946_c0_g1         ------------------------------------------------------------     734
Gvel_16103_c0_g1         ------------------------------------------------------------     504
Gvel_83898_c0_g1         CGAGGCGAAGCTGAAGGAGCTGCAGCGCGACTGCGCGCCCGTCATGACCAAGCTGCACCA     603
                         :***:..::....:**.*****..*..:*.:***.*:**::*:***.*:***********

S.americana.Hsp70A1      GGGCACTAAGGATCCCAGCAGTAGCTGTGGGCAACAAGCAGGTACAGGAGCAGCACACAC     2097
S.americana.HSP70B2      GGCGGGCGCCGGCGGCGGCGCGGGCGCAGGCACGGGCTCC------------GGGGCCGC     1929
S.serialis.HSP70B2       GGCGGGCGCCGGCGGCGGCGCGGGCGCAGGCACGGGCTCC------------GGGGCCGC     1925
Gvel_19946_c0_g1         ------------------------------------------------------------     734
Gvel_16103_c0_g1         ------------------------------------------------------------     504
Gvel_83898_c0_g1         GGGCGCGGCGGGCCAGGGCGGCGCGCC---------------------------GGGCGC     636
                         **..:..::.*::..::**:..:::.:.::........:.            :.::.*:*

S.americana.Hsp70A1      CGGTGGACCAACTATTGAGGAGGTTGACTAACTGAACTG---------------------     2136
S.americana.HSP70B2      CAAGGGCCCCACCGTCGAGGAGGTCGACTGAGGCGGCGGCGAGGCCGCTCCGTGAAGAAC     1989
S.serialis.HSP70B2       CAAGGGCCCCACCGTCGAGGAGGTCGACTGAGGCGGCGGCGAGGCCGCTCCGTGAAGAAC     1985
Gvel_19946_c0_g1         ------------------------------------------------------------     734
Gvel_16103_c0_g1         ------------------------------------------------------------     504
Gvel_83898_c0_g1         CAAGGGCCCCACCGTGGAGGAGGTGGACTAGACGCCCCGCCCTGCCCCGCCCCGCGCCGC     696
                         *:::**:**:**::*:********.****.:.....*.*:...:::.:.::..:.....:

S.americana.Hsp70A1      --------------------------------------------------TTAATATA--     2144
S.americana.HSP70B2      TCTACTGGACTGTGGCCACACCATTCTAACACACTCCTTACACTAGCCTGCCACTCTCGT     2049
S.serialis.HSP70B2       TCTACTGGACTGTGGCCACACCATTCTAACACACTCCTTACACTAGCCTGCCACTCTCGT     2045
Gvel_19946_c0_g1         ------------------------------------------------------------     734
Gvel_16103_c0_g1         ------------------------------------------------------------     504
Gvel_83898_c0_g1         CCCCGCCCCCGCCCCCGA---------CACGCACGCGGGA--C-------GCGCCGGCGC     738
                         .:.......:.....:.:......... ::.:::.:...:..:........::::.:::.
```

**Figure S4 (8/8)**

```
S.americana.Hsp70A1      ------AAAACGGTTTTACGTGTCACTGTGTGTGATATATAAAATGTGGTGTACATAGCA     2198
S.americana.HSP70B2      TTGGGAATTACTTCGTTGTGCGTCACACACTATGCTCCAAATTTTGTTGTCCCCGTATTG     2109
S.serialis.HSP70B2       TTGGGAATTACTTCGTTGTGCGTCACACACTATGCTCCAGATTTTGTTGTCCCGGTATTG     2105
Gvel_19946_c0_g1         ------------------------------------------------------------     734
Gvel_16103_c0_g1         ------------------------------------------------------------     504
Gvel_83898_c0_g1         TCGCGAAC----------------------------------------------------     746
                         :.:.::*..::.....::...:.:::::.....:.::.:...: :...:::.::.......::...
```

**Figure S4 (above)**. Partial alignment of putative *Gryllus veletis* HSP70 transcripts GGSD01027946.1; (Gvel_19946_c0_g1), GGSD01022798.1 (Gvel_16103_c0_g1) and GGSD01130040.1 (Gvel_83898_c0_g1_i1) against other orthopteran HSP nucleotide sequences from NCBI: XM_047139358.1 (*Schistocerca americana* Hsp70-A1), XM_047140202.1 (*S. americana* Hsp70-B2), and XM_050106196.1 (*S. serialis* Hsp70B2). Produced using Clustal Omega, and modified with the following symbols: one dot (.) indicates the same nucleotide in two sequences, two dots (:) indicates the same nucleotide in two sequences, and an asterisk (*) indicates the same nucleotide in four sequences.
